# Voice and Face Gender Perception engages multimodal integration via multiple feedback pathways

**DOI:** 10.1101/2020.01.07.884668

**Authors:** Clement Abbatecola, Kim Beneyton, Peggy Gerardin, Henry Kennedy, Kenneth Knoblauch

## Abstract

Multimodal integration provides an ideal framework for investigating top-down influences in perceptual integration. Here, we investigate mechanisms and functional networks participating in face-voice multimodal integration during gender perception by using complementary behavioral (Maximum Likelihood Conjoint Measurement) and brain imaging (Dynamic Causal Modeling of fMRI data) techniques. Thirty-six subjects were instructed to judge pairs of face-voice stimuli either according to the gender of the face (face task), the voice (voice task) or the stimulus (stimulus task; no specific modality instruction given). Face and voice contributions to the tasks were not independent, as both modalities significantly contributed to all tasks. The top-down influences in each task could be modeled as a differential weighting of the contributions of each modality with an asymmetry in favor of the auditory modality in terms of magnitude of the effect. Additionally, we observed two independent interaction effects in the decision process that reflect both the coherence of the gender information across modalities and the magnitude of the gender difference from neutral. In a second experiment we investigated with functional MRI the modulation of effective connectivity between the Fusiform Face Area (FFA) and the Temporal Voice Area (TVA), two cortical areas implicated in face and voice processing. Twelve participants were presented with multimodal face-voice stimuli and instructed to attend either to face, voice or any gender information. We found specific changes in effective connectivity between these areas in the same conditions that generated behavioral interactions. Taken together, we interpret these results as converging evidence supporting the existence of multiple parallel hierarchical systems in multi-modal integration.

## Introduction

Hierarchy has long played a prominent role in models of cortical organization and processing (Felleman & Van Essen, 1991; Ferster et al., 1996; Hubel & Wiesel, 1962, 1965). For example, understanding receptive field structure via the elaboration of visual information in the ascending visual pathways has been a highly successful approach to dissecting cortical physiology (J. Freeman et al., 2013). Feedforward models of receptive field construction are usefully informed by the integration of structure and function in large-scale laminar models of feedback/feedforward processes of cortical information flow (Baldauf & Desimone, 2014; Kravitz et al., 2013; Markov et al., 2014). The relevance of connectivity based structural hierarchy (Markov et al., 2014) has been recently reinforced by its remarkable agreement with functional hierarchy derived from inter-areal causal relations based on synchronization of different frequency bands of the local field potential obtained with surface electrodes in macaque (Bastos et al., 2015) and magnetoencephalography recordings in human (Michalareas et al., 2016).

While feedforward processes are typically equated with bottom-up processes, such as stimulus coding and feature selectivity, the role of feedback is more elusive and has been studied in relation to several top-down functions, including attention, perceptual integration, prediction and mental imagery. This problem is further complicated by the existence of complex networks that cannot be described uniquely in terms of a single pair of feedback/feedforward communication channels (Bergmann et al., 2019). Multiple functional pathways are, in fact, supported anatomically, an example of such being the dual counterstream architecture (Markov & Kennedy, 2013).

In this context, multi-modal integration as a model system can be highly informative. For example, auditory activity has been reported to have a predictive role in the primary visual cortex (Petro et al., 2017). In the present report we examine the multimodal integration of vision and audition as a high level feature of voice and face perception as evidenced in gender perception. In both the visual and auditory systems, information from both modalities provides a readily identifiable top-down cross modal signal. Hence, multimodal integration via a psychophysical paradigm has permitted us to evaluate models of top-down and bottom-up information integration. Finally, in order to localize neural correlates of the perceptual integration process we employ functional imagery to elucidate the effective connectivity underlying the behavioral responses.

### Multimodal face-voice integration as a window to hierarchy

Because face and voice stimuli conjointly allow the retrieval of similar information about others (e.g., identity, health, age, emotional state etc.), it has been proposed that they share a privileged link (Belin, 2017; Campanella & Belin, 2007; Yovel & Belin, 2013). Infants as young as two months match lip movements and vocal production (Patterson & Werker, 2003). In adults, distinctive voices facilitate the recognition of unfamiliar faces compared to non-distinctive voices, but there is no similar effect of distinctiveness for other auditory stimuli (Bülthoff & Newell, 2015). Simultaneous presentation of a face and a voice (but not a voice with a non-face or a non-voice with a non-face) facilitates auditory recognition (von Kriegstein & Giraud, 2006). Complementary adaptation effects within each modality suggest similar norm-based perceptual coding relative to a prototypical “neutral” face or voice, although the existence of cross-modal adaptation is controversial (Little et al., 2013; Schweinberger et al., 2008).

Despite originating from different sensory inputs, face and voice perception engage similar brain mechanisms, facilitating their multimodal integration (Belin, 2017; Campanella & Belin, 2007; Hasan et al., 2016; Yovel & Belin, 2013). In particular the Fusiform Face Area (FFA), a functionally defined region in the temporal fusiform gyrus, has been shown to respond strongly to faces (Kanwisher et al., 1997; Kanwisher & Yovel, 2006), while the Temporal Voice Area (TVA), a region in the lateral temporal cortex, has been proposed to be the equivalent of the FFA for voices (Belin et al., 2000; Pernet et al., 2015; von Kriegstein & Giraud, 2004). These areas are closely linked; Schall et al. (2013) reported responses in the FFA 110ms after voice onset. There are strong parallels in the two pathways. Lesions in the face stream induce prosopagnosia, a face-specific deficit (Davies-Thompson et al., 2014) while lesions in the voice stream induce phonagnosia, a voice-specific deficits (Roswandowitz et al., 2018). Although face-specific and voice-specific deficits do not always occur together (Arnott et al., 2008; Van Lancker & Canter, 1982), there exist several multimodal interaction effects that, however, are not always symmetrical. Hence, contrary to control subjects, voice recognition in prosopagnosics does not benefit from bimodal learning, while voice recognition in phonagnosics does benefit from bimodal learning (Maguinness & von Kriegstein, 2017). The TVA also responds to face stimuli in early deafness (Benetti et al., 2017) although the reverse is not the case for voice responses in the FFA of early blindness (Dormal et al., 2018).

Face-voice multimodal integration phenomena have been observed across species. Rhesus macaques are able to associate the face with the voice of familiar individuals, both for conspecifics and humans (Habbershon et al., 2013; Sliwa et al., 2011), and they successfully recognize vocalizations with associated facial expressions in conspecifics (Ghazanfar & Logothetis, 2003). Macaques are also able to combine noisy face and voice information so as to enhance their detection of vocalizations, a task on which they perform comparably to humans (Chandrasekaran et al., 2011). Invasive studies in non-human primates reveal an unsuspected complexity in the primate representation of voices and faces. In the macaque, for example, face responsive cortex comparable to the FFA has been found in the peripheral middle-lateral temporal lobe (Lafer-Sousa & Conway, 2013) and voice responsive neurons in the parabelt region (Perrodin et al., 2011), where it has been argued they form a TVA-like area (Belin et al., 2018). Multimodal neurons reacting to bimodal (face and voice) communication have been also discovered in the auditory cortex, the superior temporal sulcus and frontal areas (Ghazanfar et al., 2008; Ghazanfar et al., 2005; Sugihara et al., 2006). Neurons reacting to stimulus incoherence (faces with incongruent vocalizations) were found in the prefrontal cortex (Diehl & Romanski, 2014). Concerning this last study in particular, it should be noted that no other areas were tested so these results do not exclude the existence of incongruence effects elsewhere in the brain.

In studying the neural mechanisms of face and voice perception in human, there are numerous attributes that can be analyzed: identity, emotion, gender, etc. In the current study, we focus on gender perception as it presents several advantages. Face-voice gender recognition is robust and precocious, appearing as early as 6-8 months in human development (Patterson & Werker, 2002; Walker-Andrews et al., 1991). Face (and presumably voice) gender discrimination has been reported to be relatively costless in terms of attentional resources (Reddy et al., 2004). Importantly from a psychophysical point of view, and contrary to other face-voice properties such as emotions, gender is defined along a single perceptual dimension (varying between masculine and feminine) simplifying analyses of responses. Technically, it is possible to generate a continuous physical variation between male and female stimuli through morphing of both sound and image stimuli.

There is evidence that TVA and FFA are both involved in the unimodal recognition of gender in their respective modalities (Charest et al., 2012; Contreras et al., 2013; Kaul et al., 2011; Weston et al., 2015). In addition, both of these areas along with a supramodal fronto-parietal network have been found to be activated during a face-voice gender categorization with a functional magnetic resonance imaging (fMRI) protocol (Joassin et al., 2011). Patterns of activation that discriminate gender in faces (and also facial expressions) have been reported in early retinotopic areas (Petro et al., 2013), a finding that raises interesting possibilities with respect to feedback processes. Behaviorally, gender categorization of faces can be perturbed by the simultaneous presentation of pure tones in the male or female fundamental-speaking-frequency range (Smith et al., 2007). This result has been reproduced with voices instead of non-voice-like tones and shown to be a continuous process during decision making (J.B. Freeman & Ambady, 2011). While this effectively demonstrates the disruptive effect of voice gender during face gender recognition it does not give access to the underlying neural mechanisms. A mechanistic approach requires adopting a symmetrical paradigm involving simultaneous presentation of faces and voices both varying in gender, thereby allowing the collection of gender evaluations for the two dimensions.

Two studies previously used such a paradigm (Latinus et al., 2010; Watson et al., 2013) and reported mutual effects of gender between the two modalities. In a gender identification task of natural faces and voices, Latinus et al. (2010) observed that incongruent face information reduced the proportion of correct gender categorizations of voices but incongruent voice information did not reduce gender categorization of faces. They argued that gender information is more easily and rapidly extracted from faces than voices and that this extraction is more automatic, perturbing the direction of attention to voices. In contrast, Watson et al. (2013) used audio and video morphing to obtain a more fine-grained and controlled gradation of gender in both modalities. They observed that voices were more disruptive for face judgments than were faces for voices, which they attributed to the greater dimorphism of voices compared to faces with respect to gender.

Both of the above studies used a single-stimulus gender identification paradigm in which they measured the probability of a male/female decision. It is difficult to compare face and voice contributions on a common scale by simply comparing probabilities. Instead, it would be preferable to model the common underlying decision process. Additionally, while Watson et al. (2013) could detect cross-modal effects, they failed to describe such interactions qualitatively and quantitatively. In the current study, we use the psychophysical method of Maximum Likelihood Conjoint Measurement (MLCM) that obviates these problems by evaluating gender comparisons within a signal detection model of the decision process. This approach formalizes a set of nested models describing several types of face-voice integration that can be tested (Ho et al., 2008; Knoblauch & Maloney, 2012). Having characterized the nature of the multi-modal interactions behaviorally, we employed a functional imaging paradigm to identify inter-areal interactions that follow similar patterns of face-voice integration.

### Maximum Likelihood Conjoint Measurement

MLCM is a scaling method in which paired-comparison judgments are made between stimuli that co-vary simultaneously and independently along *n* ≥ 2 stimulus dimensions (Ho et al., 2008; Knoblauch & Maloney, 2012). MLCM has been used to investigate a diverse set of perceptual processes, including visual surface perception (Hansmann-Roth & Mamassian, 2017; Ho et al., 2008; Qi et al., 2015), stimulus dependence in a visual filling-in illusion (Gerardin et al., 2014; Gerardin et al., 2018), time constancy in perception (Lisi & Gorea, 2016), lightness and chroma perception in adults (Rogers et al., 2016) and infants (Rogers et al., 2018), and perception of light scattering and diffusion in liquids (Chadwick et al., 2018). In the present case, the stimuli are defined as variations in the face and voice gender along a continuum of interpolated faces and voices obtained by morphing the extreme exemplars. The contributions of each dimension and their interaction are estimated via a signal detection model of the decision process that is fitted by maximum likelihood to best predict the observer’s choices over the set of all trials. These contributions quantify the extent to which judgments along one dimension (face or voice) depend on the morphing levels of the other (voice and face).

Three nested models describing the contribution of the dimensions tested to the decision process are fit to the data and evaluated with likelihood ratio tests: an Independence model in which only one dimension (e.g. the face morphing level while judging face gender) contributes to the judgments; an additive model in which the contribution of each dimension depends only on its level along the dimension and judgments are based on the additive sum of each contribution; and finally a saturated (full or interaction) model where judgments depend on the particular combination of stimulus levels on each dimension.

In MLCM models, non-additive combination terms are usually introduced as non-specific interaction effects (Knoblauch & Maloney, 2012), but we can instead parameterize them according to a priori hypotheses. To motivate such hypotheses, we consider the combination of multi-modal information within the context of optimal cue combination, i.e., how does the observer combine or integrate two independent cues to arrive at a decision.

### Signal Detection Theory and optimal cue integration

Clark and Yuille (2013) defined two classes of models to describe how several cues can be fused into a single percept. In weak fusion models each cue is first processed separately in distinct and independent modules, after which a rule is used to combine them. By contrast, strong fusion models de-emphasize modularity and allow interaction effects at any processing level. Young et al. (1993) and (Landy et al., 1995) established the modified weak observer model as a variant of weak fusion that includes: i) early interactions to make cues commensurable, whereby psychological variables, *ψ*_V_ and *ψ*_A_ along dimensions subscripted as V and A for visual and auditory, respectively, are expressed in the same units on the same internal perceptual axis and can, thus, be directly added, ii) the possibility of other potential interaction effects when the discrepancy between individual cues is beyond that typically found in natural scenes, e.g., constraining physical values, *ϕ*_V_ and *ϕ*_A_, such that *ψ*_V_ ≈ *ψ*_A_ to prevent such effects and iii) a weighted average cue combination rule with weights dynamically chosen to minimize variance of the final estimate. This leads to the following signal detection model of the psychological response:

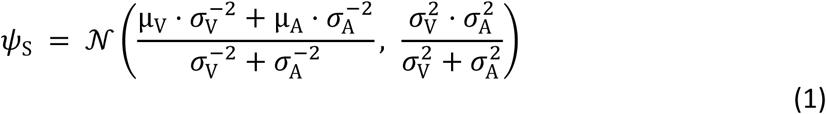

where *ψ*_S_ is the perceived level of the fused stimulus, distributed normally with *μ*_i_ the mean perceived level of the stimulus along dimension *i*, and 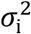 the variance of the perceived level along dimension *i* (for derivation and a fuller description of this model see Supplementary Section 4).

The consequences of this model are that increasing either variance brings the variance and the mean of the combined percept closer to whichever estimate is more reliable. The combined percept is always expected to be more precise than either isolated unimodal estimate. Note that the observer is predicted to make binary decisions about combined stimuli, that is, classify them as either feminine or masculine in the current case, in proportion to the position of the density distribution along the perceived gender axis with respect to a gender-neutral value.

Observers who use this maximization rule with respect to reliability are called statistically optimal and this type of response has been empirically verified in several domains for human multimodal integration (Alais & Burr, 2004; Ernst & Banks, 2002; Hartcher-O’Brien et al., 2014).

As already mentioned, one limitation of this model is its dependency on the condition *ψ*_V_ ≈ *ψ*_A_. In other words complementary interaction effects are expected to come into play in the case of inconsistencies beyond the discrimination threshold between the cues. In the absence of such interaction effects, the resulting percept is predicted to be the same when combining visual and auditory cues as long as the variances and 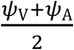 remain constant.

An extension of the model includes an interaction effect of coherence. When, for example, one modality is very masculine and the other very feminine, we make the hypothesis that the overall stimulus will be perceived as less reliable than in a more coherent context. Consequently the variance of each unimodal estimate is increased in proportion to the distance between them in gender space before applying the optimal integration scheme. This has the effect of lowering also the variance of the combined gender estimate, which biases the decisions because a higher proportion of the distribution falls on one side of the gender neutral reference.

Beyond this effect of coherence, it is also conceivable that there is an effect of the magnitude of the gender difference from neutrality contained in each modality. For example, a face that is very clearly masculine might generate a more precise representation than a more gender-neutral face. Such effects would become more significant with greater incoherence between cues due to an increase in the difference in precision, and once again the resulting change in variance of the final estimate biases decisions.

The above models can be implemented by introducing an internal coherence parameter and/or a multiplicative effect of gender to take into account the quality of gender information.

### Dynamic Causal Modeling

While the psychophysical results can suggest the nature of the interactions between modalities, they do not localize sites and pathways that participate in the phenomena under study. To explore the neural substrate of the multi-modal integration revealed by the MLCM experiments, we performed a series of functional imaging experiments in which we evaluated the effective connectivity between areas implicated in processing face and voice stimuli. In particular, we restricted our consideration to two candidate areas, the FFA (Kanwisher et al., 1997; Kanwisher & Yovel, 2006) and the TVA (Belin et al., 2000; Pernet et al., 2015; von Kriegstein & Giraud, 2004). The FFA and TVA are ostensibly unimodal modules involved, respectively, in face and voice processing. We translated the MLCM models described above into hypothetical effective connectivity networks in order to test whether non-additive interactions involve unimodal areas or occur exclusively at higher levels. Importantly, observing changes in effective connectivity between the FFA and TVA is agnostic to the role of direct communication between these areas or mediated by other top-down effects.

Figure 1 presents a series of schematic networks of multimodal integration based on the type of models of inter-areal connectivity that best describes different potential results of the MLCM experiments. The independence model would implicate direct communication between unimodal face and voice areas and the site of gender decision, whereas its rejection would imply the existence of at least one mandatory site of multimodal integration prior to the gender decision. Models with interactions are more challenging to interpret. The substrate of interaction effects is presumably a change in connectivity in an inter-areal network. There are two possibilities: either the unimodal areas are themselves involved in this interaction, or the changes occur exclusively at designated multimodal sites, presumably at levels hierarchically above the unimodal areas. It is not possible to decide between these two hypotheses using psychophysical experiments alone, but the question can be addressed by looking at brain activity obtained from fMRI. Dynamic Causal Modeling (DCM) with Bayesian Model Selection (BMS) (Penny, 2012; Penny et al., 2010) permits evaluation of the capacity of activity in one brain area to cause or generate activity in another (effective connectivity). If such changes of effective connectivity are observed under the same conditions that we observe behavioral interaction effects, it is possible to conclude (1) that these changes are indeed a neural correlate of interaction and (2) that unimodal areas are involved.

**Figure 1:**
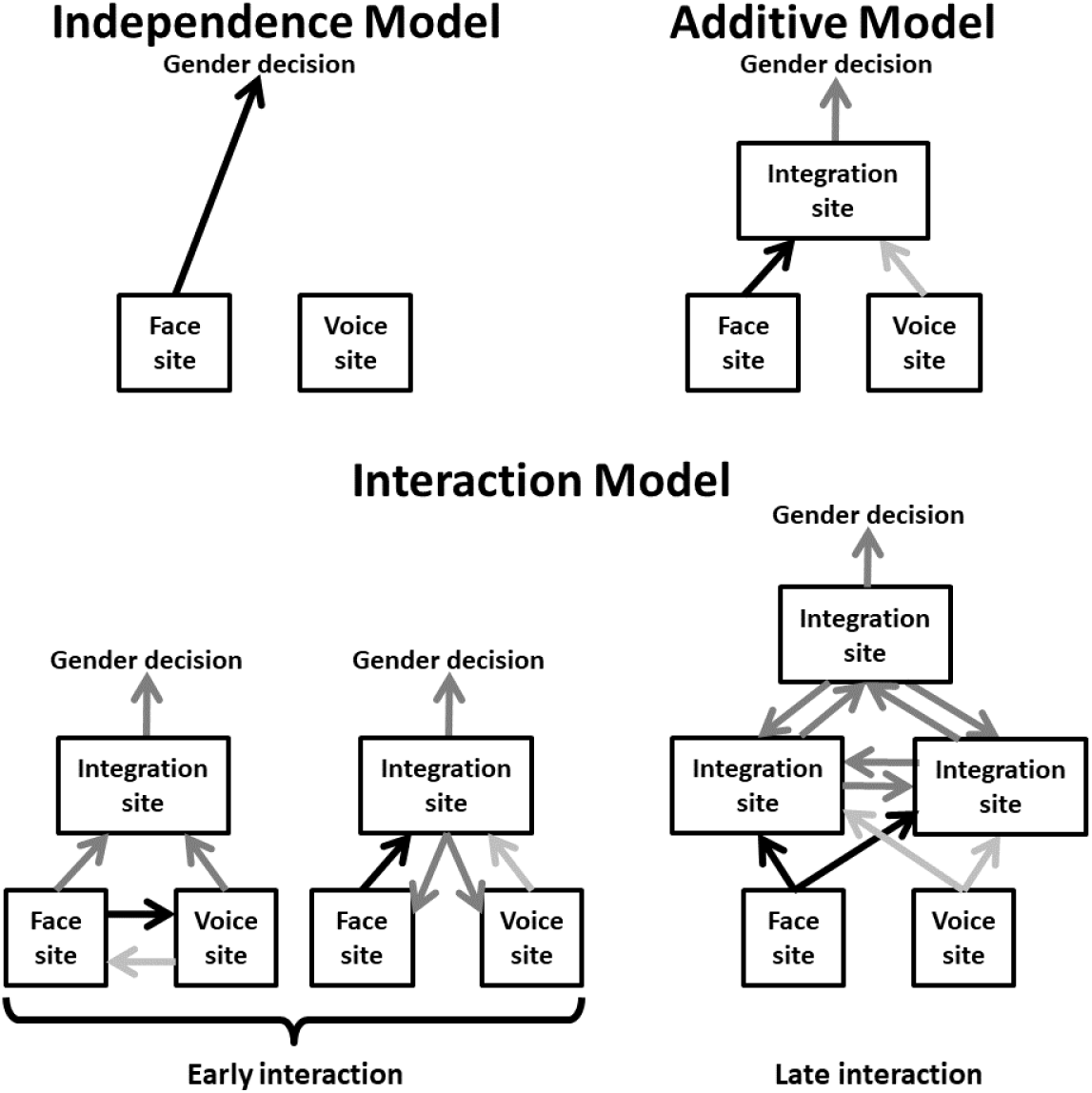
Underlying cerebral processes suggested by MLCM models for face-voice gender integration. Top left: the independence model suggests a direct link between unimodal face and voice sites and the site of gender decision. Top right: the additive model implies the existence of at least one mandatory site of multimodal integration before gender decision. Bottom: the saturated model is compatible with several interpretations that can be divided in two groups depending on whether the non-additive interaction involves unimodal areas or takes place exclusively at a higher level.

## Material and methods

### Psychophysics

#### Observers

Thirty-six observers (18 male) with normal or corrected-to-normal vision volunteered for the experiment (mean age +/- SD: 25.9 +/- 3.6 years). Each was randomly assigned to one of the six experimental conditions so that there were 3 male and 3 female observers per condition. All observers but one (author PG) were naive, all were right handed, native French speakers. All observers had normal (or corrected to normal) vision as assessed by the Freiburg Visual Acuity and Contrast Test (FrACT) (Bach, 2007), and the Farnsworth F2 plate observed under daylight fluorescent illumination (Naval Submarine Medical Research Laboratory, Groton, CT, USA). Normality of face perception was assessed by the Cambridge Face Memory Test (CFMT) (Duchaine & Nakayama, 2006). All observers gave informed consent.

#### Apparatus

The experiments were performed in a dark room. Stimuli were displayed on an Eizo FlexScan T562-T color monitor (42 cm) driven by a Power Mac G5 (3gHz) with screen resolution 832 × 624 pixels and run at a field rate of 120 Hz, non-interlaced. Calibration of the screen was performed with a Minolta CS-100 Chromameter. Observers were placed at a distance of 57.3 cm from the screen. Head stabilization was obtained with a chin and forehead rest. Auditory stimuli were presented through headphones (Sennheiser HD 449), which also served to mask any ambient noise. Sound calibration was performed with a Quest QE4170 microphone and a SoundPro SE/DL sound level meter.

#### Stimuli

The stimulus set, obtained from Watson et al. (2013), consisted of video clips of a person saying the phoneme “had”, whose face and voice varied by morphing from feminine to masculine (18 levels of morphing for the face and 19 levels for the voice yielding a total of 342 combinations). The clips were converted to greyscale and matched for luminance. An oval mask fitted around each face hid non-facial gender cues, such as the hair and the hairline (see Figure 2 for examples).

**Figure 2:**
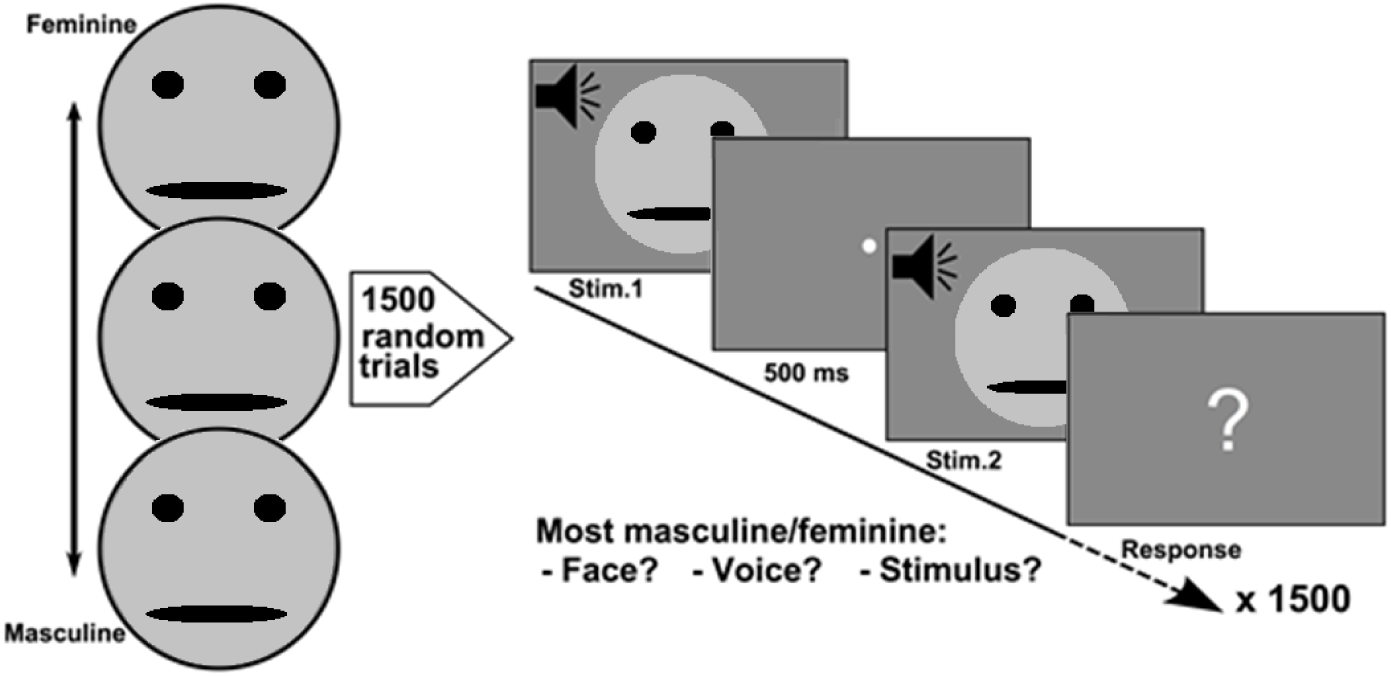
Stimulus set and Conjoint measurement protocol. Pairs of face-voice video sequences with independently varying levels of face and voice gender morphing were compared by the observers according to: (1) face gender, (2) voice gender or (3) stimulus gender taking both face and voice into account. Stimulus pairs were sampled from a set with 18 levels of morphing for the face and 19 levels for the voice. Six groups of observers judged, respectively, which face, voice or stimulus was either most masculine or feminine (6 observers/group, each group balanced with respect to gender, 36 observers total, 1500 trials/observer). Note that faces were replaced by cartoons for the bioRxiv manuscript.

The software PsychoPy v1.80.07 (Peirce, 2008) was used to control stimulus presentation. Stimuli were displayed in the center of a grey background (31.2 cd/m^2^, CIE *xy* = 0.306, 0.33). Face luminance varied between 29.7 cd/m^2^ (CIE *xy* = 0.306, 0.324) for the eyes and 51.6 cd/m^2^ (CIE *xy* = 0.303, 0.326) for the nose. Face diameter was fixed at 10 cm and voice volume between 85.2 and 86.7 dB SPL (A) - Peak.

#### Procedure

Observers were tested over five sessions of 300 trials each, yielding a total of 1500 trials per observer (2.57% random subsampling from the ((19 × 18 – (19 × 18 – 1))/2 = 58311 total number of non-matching, unordered pairs). Given the large number of possible pairs tested, a subsampling paradigm was necessary to make the number of trials that each subject was tested tractable. As this is a novel approach for performing MLCM experimentally, simulations justifying this procedure are presented in Supplementary Section 1. On each trial two stimuli were randomly selected with the gender morphing scale values of the voice and face independently and randomly assigned and successively presented (Figure 2). The duration of each stimulus was fixed at 500ms with a 500ms inter-stimulus interval between each pair. After the presentation, observers were prompted to make a judgment comparing the genders of the two stimuli. The next pair was presented following the observer’s response via a button press. Observers were randomly assigned to one of six groups with the constraint that each group was composed of equal numbers of males and females. The groups were defined by whether the observers were instructed to judge on the basis of the face, the voice or the stimulus, i.e., no specific instruction regarding modality, and whether they were instructed to choose which of the pair was more masculine or feminine. Six observers were tested in each of the 6 conditions.

#### Analysis

MLCM aims to model the decision process of observers comparing multidimensional stimuli to determine how the observer integrates information across dimensions to render a judgment. Because the decision process is noisy, a signal detection framework is used (Ho et al., 2008) and the resulting model can be formalized as a binomial Generalized Linear Model (Knoblauch & Maloney, 2012). Several nested models, corresponding to increasingly complex decision rules for combining the information across dimensions, are fitted to the data using maximum likelihood in such a way as to maximize the correspondence between model predictions and observer decisions. These models are compared using nested likelihood ratio tests to determine the degree of complexity required to describe the observer’s decisions.

Consider two face-voice stimuli defined by their physical levels of morphing for visual and auditory gender, 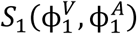 and 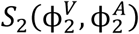, and the task of deciding whether the first or second stimulus has the most masculine face, i.e., the visual task. The noisy decision process is modeled as:

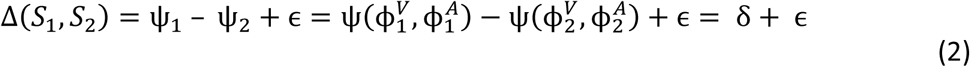

Where Ψ_1_ and Ψ_2_ are internal representations for the gender of the first and second face, respectively determined by the psychophysical function Ψ and ϵ is a Gaussian random variable with mean μ = 0 and variance σ^2^. The log-likelihood of the model over all trials given the observer’s responses is given by:

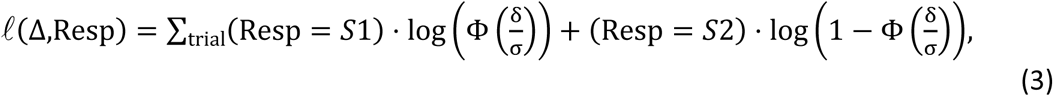

where (Resp = Si) = 1 when true, otherwise 0. Under the independence model the observer exclusively relies on the visual information. In that case we can define 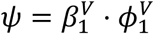 and:

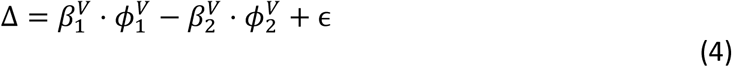

Under the additive model both visual and auditory information are combined additively. In that case we can define 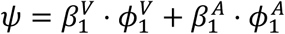 and rearranging terms:

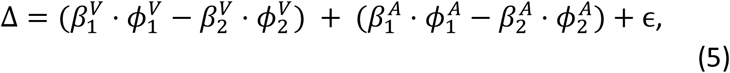

which demonstrates that, in effect, the observer is comparing perceptual intervals along one dimension to perceptual intervals along the other (Knoblauch & Maloney, 2012).

Under the interaction model, non-additive combination terms are introduced. In the present study we tested two types of interaction terms. The *Coherence Interaction Model* introduces an internal coherence effect between face and voice gender within stimulus:

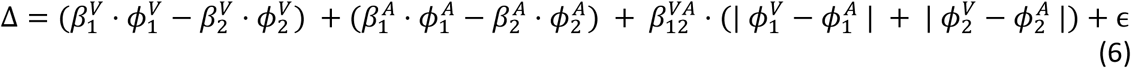

The *Magnitude Interaction Model* introduces a multiplicative effect of gender information across stimuli:

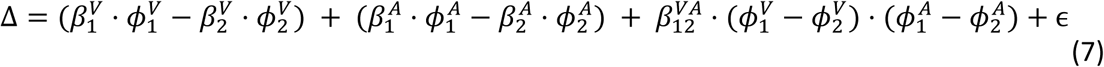

All psychophysical data were analyzed using R (R Core Team, 2019) and the package lme4 (Bates et al., 2015) to take inter-individual variability into account with generalized mixed-effects models (GLMM) (Pinheiro & Bates, 1978).

### fMRI design and procedure

#### Observers

Twelve observers (6 female, mean ± SD age: 23.3 ± 3.3 years) participated in the study. All were right handed and screened for normal vision with the same battery of tests used in experiment 1. The study was approved by the local ethics committee (ID-RCB 2015-A01018-41). In accordance with the protocol, each subject was pre-screened by a clinician for suitability to undergo an fMRI procedure and gave informed consent. All participants attended two MRI sessions of approximately 1.5 hour each and were remunerated for their participation.

#### Data Acquisition

All experiments were conducted using a MAGNETOM Prisma 3T Siemens MRI scanner at the Lyon MRI facility PRIMAGE, France. In each session for each individual, a high-resolution T1-weighted structural image (acquisition matrix 256×256×192, TR/TE: 3500/3.42ms, flip angle: 8°, resolution: .9x.9x.9mm) and series of T2*-weighted functional images (acquisition matrix 78×78, 49 slices, TR/TE: 2500/26ms, flip angle: 90°, resolution: 2.7×2.7×2.7mm) were collected with a 64 channel head/neck Siemens coil. In order to control participants wakefulness, the left eye over the course of all experiments was monitored with an SR Research EyeLink 1000 Plus. Sound was provided using NordicNeuroLab earplugs. Subject responses were recorded using a Current Designs 904 diamond shaped 4 buttons device.

#### Stimuli

The stimuli were selected from a reduced set of those used in experiment 1 (Watson et al., 2013), consisting of video clips of a person saying the phoneme “had”. We used only 3 levels of morphing for the face and 3 levels for the voice (the most feminine, gender neutral and masculine in terms of physical morphing in both cases) for a total of 9 combinations. The software PsychoPy v1.84.2 (Peirce, 2008) was used to control stimulus presentation. Stimuli were displayed in the center of a grey background with a resolution of 1024×768 pixels. The face diameter was fixed at 10 degrees of visual angle.

#### Procedure

### Face-voice protocol

Face-voice stimuli were presented in an event-related protocol. Each stimulus lasted 0.5s and was followed by a fixation point, the duration of which was randomly chosen between 5, 5.5, 6, 6.5 and 7s. To prevent habituation effects we ensured that no more than three successive repetitions occurred of the same stimulus sequence (e.g. AAA), pair of stimuli (e.g. ABABAB), and fixation point intervals.

In order to mirror the tasks of the MCLM experiment, task-dependent instructions were given to attend to the gender of the face, the voice or the stimulus (no specific instruction regarding modality), however no stimulus comparison was required. The attentional focus was controlled by randomly distributed “response” trials after which participants were prompted by a screen with a red feminine sign on one side and a blue masculine sign on the other (masculine and feminine sides varied randomly to prevent motor anticipation). They then had 2s to push a button on the left or the right side of a response device to indicate the gender perceived for the modality to which they were instructed to attend.

During each acquisition, the 9 gender combinations were presented 3 times in addition to 8 response trials composed of every combination except a neutral face and a neutral voice (which would not be informative about the attentional state of the participant), see Figure 3. All participants performed two tasks (face and voice, face and stimulus or voice and stimulus) in two sessions separated by at least one day, each session being composed of 5 acquisitions. The total number of repetitions for each condition was 15 for all face-voice combinations and 40 response trials (for which the responses, but not brain activity, were analyzed to avoid motor contamination of fMRI data). Controlling for the order of the sessions (e.g. face task first then voice task or voice task first then face task) resulted in 6 possible pairs of session, each of which was assigned to one male and one female participant. We therefore acquired 8 sessions in total for each task.

**Figure 3:**
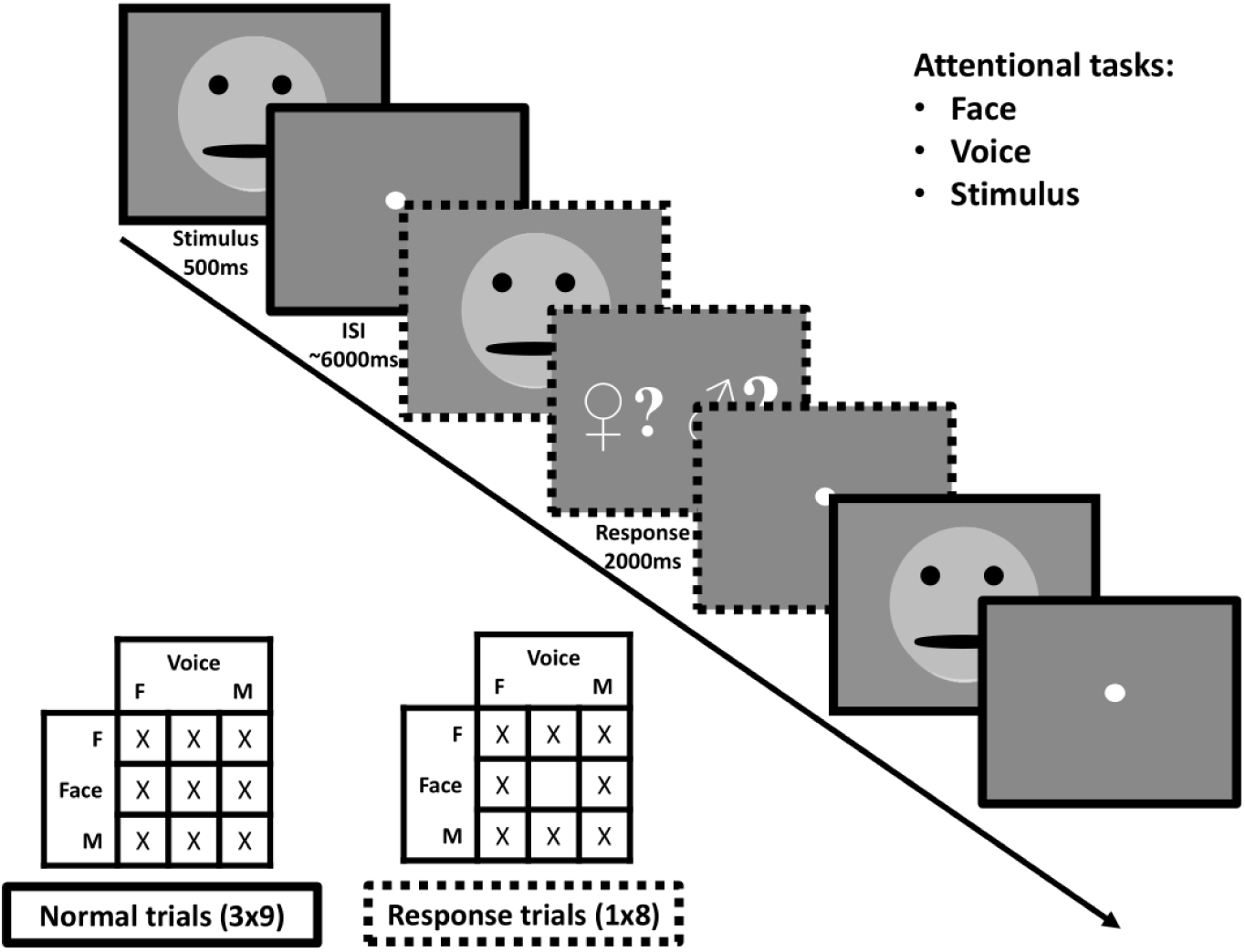
Protocol for one acquisition of the face-voice gender fMRI experiment. Top-right: within a session subjects were assigned the task to pay attention to face, voice or stimulus gender. Middle: they were presented with face-voice stimuli in succession in an event-related manner. On some trials, they received a signal to indicate whether the gender of the modality they observed was masculine or feminine for the previous stimulus (note that in the actual protocol the masculine sign was presented in blue and the feminine sign in red). Bottom-left: for each acquisition all 9 face-voice gender combinations were presented 3 times in addition to 8 “response” trials corresponding to all combinations except gender-neutral face + gender neutral voice. Note that faces were replaced by cartoons for the bioRxiv manuscript.

### Mapping regions of interest

In addition to the face-voice protocol described above we identified two regions of interest using functional localizers. During the first session we used a localizer for the FFA described by Pitcher et al. (2011) and Julian et al. (2012), in which subjects were presented with blocks of dynamic visual stimuli belonging to several categories: human faces, human body parts (hands, legs…), objects, landscapes or scrambled images (created by dividing object videos using a 15 by 15 box grid and spatially shuffling them). We reasoned that this dynamic protocol would reveal functional areas involved in our dynamic face-voice stimuli. During the second session, we used a localizer for the TVA described by Belin et al. (2000), in which subjects were presented with blocks of silence or auditory stimuli which were either vocal sounds (both with and without language) or non-vocal sounds.

#### fMRI data analysis

Imaging data were first analyzed with Brain Voyager QX (Goebel, 2012). Preprocessing functional data consisted of slice-scan time correction, head movement correction, temporal high-pass filtering (2 cycles) and linear trend removal. Individual functional images were aligned to each corresponding anatomical image. These were then used for 3D cortex reconstruction and inflation. No spatial smoothing was applied.

### Mapping Regions of Interest

FFA and TVA were identified using a General Linear Model analysis including fixation periods and movement correction parameters as nuisance covariates. The FFA was defined, bilaterally, as the set of contiguous voxels in the temporal fusiform gyrus that showed the highest activation for faces compared to body parts, landscapes, objects and scrambled images (Julian et al., 2012; Pitcher et al., 2011). The TVA was defined, bilaterally, as the set of contiguous voxels in the lateral temporal cortex that showed the highest activation for vocal compared to non-vocal sounds (Belin et al., 2000). In both cases we used a significance threshold of p<0.05 after false discovery rate correction. There is no reason to assume these functional areas are the same size across subjects but we checked that there were no outliers in terms of number of voxels (defined as a localized area that deviated from the mean number of voxels by greater than two standard deviations for an individual subject compared to the others).

### Dynamic Causal Modeling

A DCM analysis was performed using Matlab (R2014a) with SPM12 (6906). Each model was defined using the two regions of interest, FFA and TVA. Visual signals were modeled as input to the FFA and auditory to the TVA. Given the automatic nature of face-voice integration, we considered all possible connections between and within areas in the endogenous A matrix (see Figure 4, top left). In the B matrix describing effective connectivity modulations, two model spaces were defined corresponding to the two tested interaction effects:

**Figure 4:**
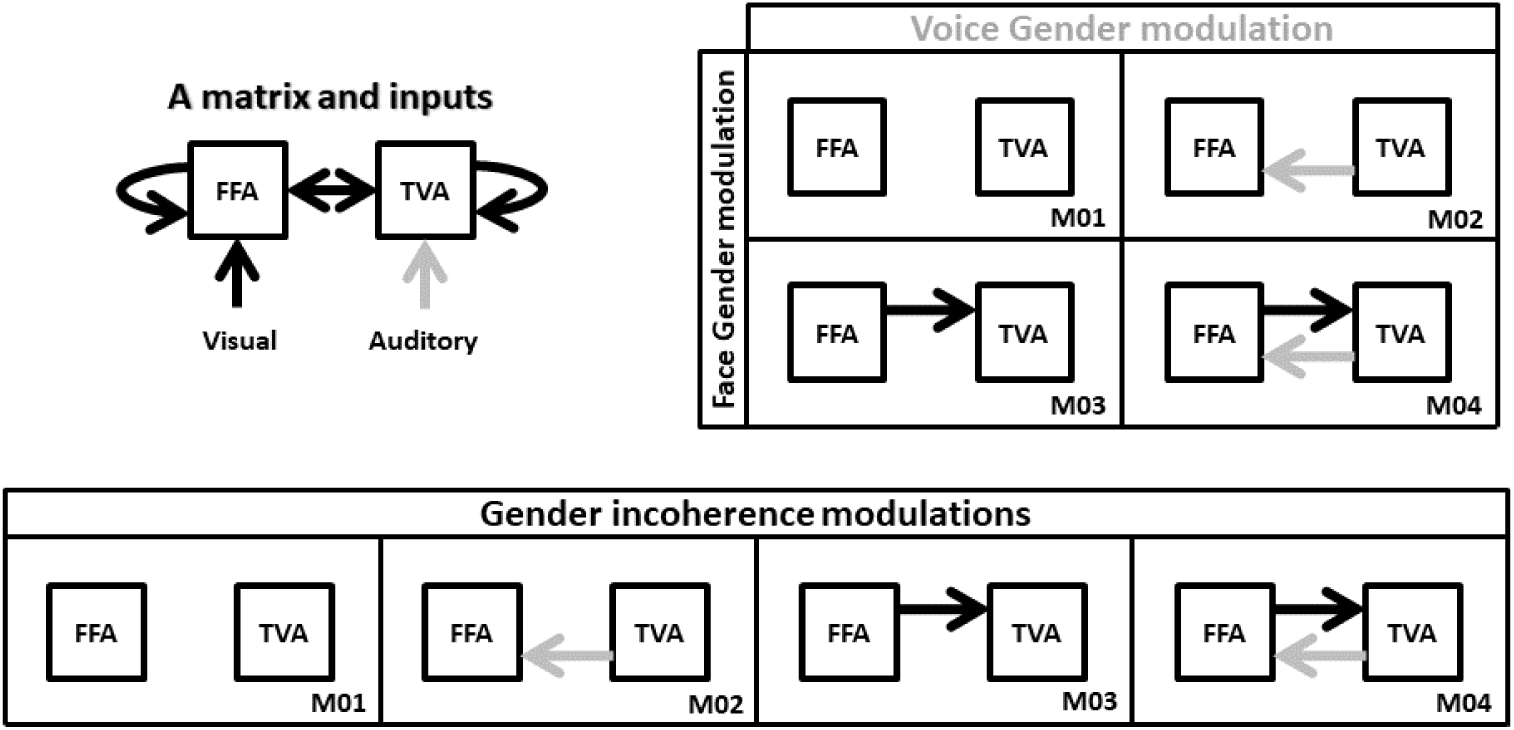
Models of DCM analysis. Top left: inputs and intrinsic connectivity that were applied to all cases. Top right: model space for changes in effective connectivity to test gender effects. Black arrows represent changes in connectivity from FFA to TVA in response to face gender (compared to gender-neutral). Gray arrows represent changes in connectivity from TVA to FFA in response to voice gender (compared to gender-neutral). Bottom: model space for changes in effective connectivity to test coherence effects. Black and gray arrows both represent changes in effective connectivity when face and voice gender are incongruent (compared to congruent).

1. Connections from FFA to TVA could either be modulated or not by added face gender information (masculine or feminine as opposed to gender-neutral); and connections from TVA to FFA could either be modulated or not by added voice gender information (masculine or feminine as opposed to gender-neutral). This results in the model space described in Figure 4, top right, with 4 possible models of modulation.
2. Compared to a gender coherent stimulus, connections between FFA and TVA could be modulated in either or both directions by a gender incongruent stimulus (i.e. a masculine face with a feminine voice or a feminine face with a masculine voice). This results in the model space described in Figure 4, bottom, with 4 possible models of modulations.

Note that, despite their apparent similarity, the two model spaces are very different in terms of the data contrasts. For example, when testing face gender modulation (i.e. models with a red arrow versus models without a black arrow in Figure 4, top-right), activation in response to the 3 stimuli with a gender neutral face (and varying voice gender) is contrasted with activation in response to the 6 stimuli with either a masculine or a feminine face (and varying voice gender). When testing coherence gender modulation from the FFA to the TVA (i.e. models with a red arrow versus models without a black arrow in Figure 4, bottom), activation in response to the 2 incoherent stimuli (with either a masculine face and a feminine voice or a feminine face and a masculine voice) is contrasted with activation in response to the 7 stimuli that are either completely coherent or gender-neutral for at least one modality.

For both model spaces, the 4 models were applied using the principal eigenvariate of the combined activation of every voxel within the FFA and the TVA of each subject. For each model within a condition (face, voice or stimulus), model evidence, i.e. the probability of observing the measured data given a specific model, was computed based on the free energy approximation using a Random-effects (RFX) analysis to account for between-subject variability (Stephan et al., 2009).

However we were not interested in the probabilities of the models per se but in the presence or absence of effective connectivity modulation from the FFA to the TVA (or vice versa) while modulation in the reverse direction is controlled. Hence, we performed family comparisons, following the procedure introduced by Penny et al. (2010). First we compared a family composed of the two models in which there is no modulation from FFA to TVA to a family composed of the two models in which there is modulation (models 1 and 2 versus models 3 and 4 from Figure 4); second we compared a family composed of the two models in which there is no modulation from TVA to FFA to a family composed of the two models in which there is modulation (models 1 and 3 versus models 2 and 4 from Figure 4). These partitions were used during the Bayesian Model Selection procedure to rank the families using exceedance probability, i.e. the probability that a family is more likely than the others in a partition, given the group data. Following previous work (Penny et al., 2010), we used a probability of .9 as strong evidence in favor of a family compared to the other but note that thresholds are less critical in this context than in a frequentist framework.

## Results

### Psychophysics

The face and voice contributions for each observer are estimated initially with the additive MLCM model and are displayed in Supplementary Figures S4 to S6. No differences were observed between male and female subjects except for the voice task (linear mixed-effects model, face task: *χ*^2^(37) = 30.7, p = 0.76; voice task: *χ*^2^(37) = 77.3, p = 0.0001; Stimulus task: *χ*^2^(37)= 49.2, p = 0.09). While this suggests a significant difference between male and female observers in how gender information is integrated while judging voices, the graph of the estimated components suggests that the effect is small and generated by a slight tendency of female observers to be more influenced than male observers by facial information for strongly male faces (Supplementary Figure S7). No differences were observed between groups that judged “more masculine” or “more feminine”, after the response reversal was taken into account (linear mixed-effects model, face task: *χ*^2^(37) =26.3, p = 0.5; voice task:: *χ*^2^(37) = 36.8, p 0.48; Stimulus task: *χ*^2^(37) =33.5, p=0.63). Given these results, we combined the data sets from male and female observers who responded “more masculine” or “more feminine” for each of the three tasks in all subsequent analyses.

The graphs in Figure 5 summarize the additive model for each of the three tasks. Each point is the average from 12 observers. The two sets of points in each graph indicate the contributions that best predict the observers’ choices over all stimulus presentations of the face (light gray) and the voice (dark gray) to the decision variable as a function of the morphing level of the stimulus, varying from most feminine (left) to most masculine (right). Hence, in the face task, the predicted internal response for a stimulus with face gender level 10 and voice gender level 15 is obtained by reading-off the ordinate value of the light grey point at 10 on the abscissa and the dark grey point at 15 and summing the two values. If the same calculation is performed for a second stimulus, the two internal response estimates can be compared with the greater value determining which stimulus is predicted to be judged as more masculine in the absence of judgment noise. The estimated scale values have been parameterized so that the standard deviation of the judgment noise corresponds to 1 unit of the ordinate values. For this reason, we specify the scale as the signal detection parameter *d’* (Green & Swets, 1966; Knoblauch & Maloney, 2012).

**Figure 5:**
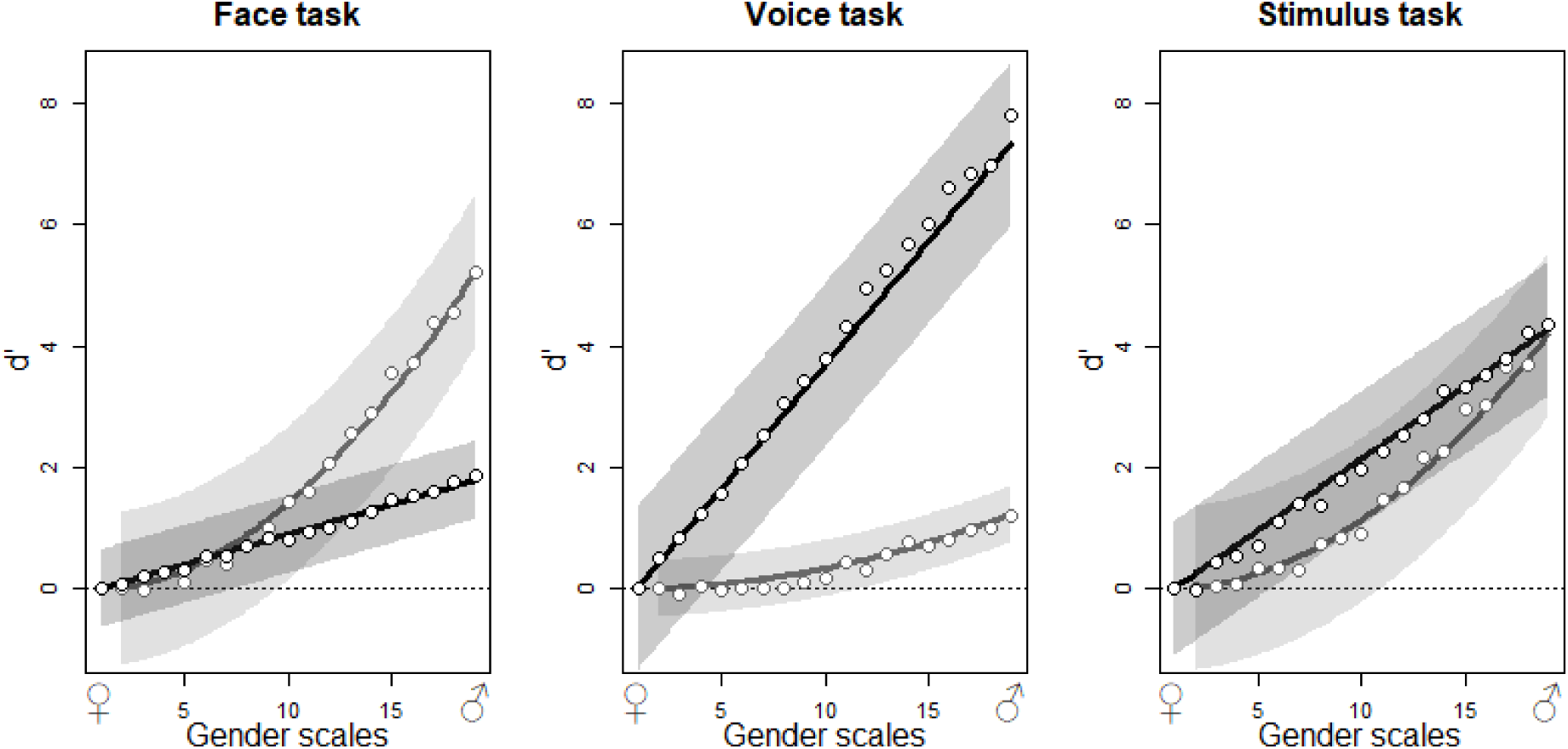
Contribution of the masculinity of the face (light gray) and the voice (dark gray) to gender decision while evaluating the gender of the face (left panel), the voice (center panel) and the stimulus (right panel). Abscissa indicates the levels of morphing of the faces and the voices from feminine to masculine and ordinates the contribution to gender judgment expressed in d’ units. Each plot represents the fixed effects and their 95% confidence interval from the additive GLMM analysis (lines and envelopes, respectively) and the mean values of observers corresponding to each level in the additive GLM analysis (points) for 12 observers (36 observers in total).

The contribution of each modality is task dependent. When observers judge the gender of the face, the face contribution is strongest with a smaller but significant contribution of the voice (Figure 5). When observers judge the gender of the voice, the contributions inverse, and when the task is to judge the gender of the stimulus, both modalities contribute about equally. In brief, effects of the task in this experiment can be conceived as relative changes of the weights of modality contributions to favor the contribution of the relevant modality.

Interestingly, the figures suggest that the task modulates only the relative contribution of each modality without changing its functional form. For all three tasks, the voice contribution varies approximately linearly with gender level with its slope varying across task. Similarly, the face contribution varies in a nonlinear fashion with gender level but the form remains similar across tasks. We tested the shape-invariance of the modality contributions (Figure S8), which shows the average values for the visual (left) and auditory (right) components for each of the three tasks, normalized to a common ordinate scale, and normalized shapes approximated with a non-linear least square approach. The voice contribution is well defined by a linear and the face contribution by quadratic functions. We refitted the MLCM models with these fixed curves that varied only with respect to a task specific coefficient using a Generalized Linear Mixed-Effects Model (GLMM) (Bates et al., 2015). The fitted curves with 95% confidence bands (Figure 5) provide a good description of the average data.

The random effect predictions for each individual (Figures S4 to S6) confirm that maximum variation in face and voice tasks is found for the relevant modality while the contribution of the irrelevant modality stays lower. During the stimulus task the contribution of both modalities varies. In fact the 12 observers from the stimulus task seem to fall into 3 separate groups: those that respond more like the observers in the face task, those that respond more like the observers in the voice task and those that tend to assign nearly the same weight to both modalities. This might be due to differing strategies in performing the task, or, alternatively, differing inherent sensitivities to the information from different modalities that influence gender judgments.

The additive model estimates of the contributions of each modality to the decision variable display invariant curves as a function of gender across the three tasks. This suggests that the mechanism influencing the contribution of each modality to the decision process is independent of the top-down influences that would be the case if the contribution of each modality originates prior to the top-down influence of the other modality, i.e., a purely unimodal site.

In the following analyses, we use the fits based on the GLMM parametric curves. This considerably reduces the complexity of the models. For the non-parametric independence model the number of parameters is one less than the number of gender levels, i.e., 17 for face and 18 for voice, but the use of a parametric curve reduces the number to only the one coefficient that controls the amplitude of the curve. For the additive model, the 37 parameters used to estimate the points in Figure 5 are reduced to 2, one coefficient for each curve. The saturated model would require 18 * 19 – 1 = 341 parameters but the use of the parametric curves reduces this number to only 3.

The independence model, for example, for the visual input becomes:

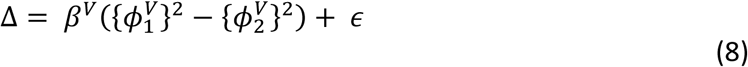

with the single parameter *β*^*V*^ and the equivalent model for an auditory input with single parameter *β*^*A*^. Similarly, the additive model becomes:

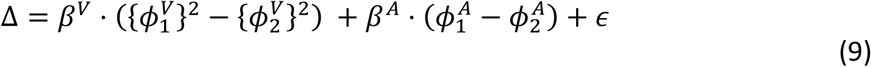

with only 2 parameters.

This framework is extended to mixed-effects model by including a random effect of observer over the coefficients. The independence (visual) model, then, becomes:

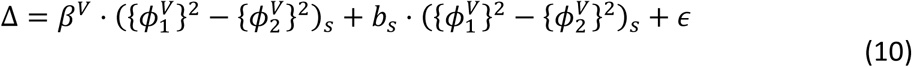

The coefficient *β*^*V*^ is a fixed effect estimate common to every subject while the *b*_*S*_ are random effect predictions assumed to be normally distributed with mean 0 and variance 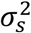. Generalization to the additive model is trivial but note that we did not include a random interaction term (only the random visual and auditory terms) for the mixed interaction models because the increased complexity of the random effects structure led to singular models, suggesting data overfitting (Bates et al., 2015).

Nested models were fit and likelihood ratio tests run to re-test the difference between male and female subject and between observers who judged the stimuli to be most masculine or most feminine based on the parametric curve estimations for each dimension. The results (Table 1) confirm previous findings suggesting that there are no differences between male and female observers for the face and stimulus tasks and only marginal evidence for a male/female observer difference in voice judgments, also supported by the change in AIC (face: *χ*^2^(2) = 1.61, p = 0.57; voice: *χ*^2^(2) = 6.6, p = 0.04; stimulus: *χ*^2^(2) = 2.25, p = 0.33). There was no evidence, however, for a difference in the fits due to the type of judgment made on any of the tasks (face: *χ*^2^(2) = 1.13, p = 0.57; voice: *χ*^2^(2) = 2.98, p = 0.23; stimulus: *χ*^2^(2) = 2.06, p = 0.36).

**Table 1:** Model comparison for psychophysical data.

| Chisq | Df | p | dAIC |
| --- | --- | --- | --- |
| **Masculine/Feminine** |  |  |  |
| 1.1312 | 2 | 0.568 | -3 |
| 2.98 | 2 | 0.23 | -1 |
| 2.06 | 2 | 0.36 | -2 |
| **Male/Female** |  |  |  |
| 1.61 | 2 | 0.45 | -2 |
| 6.60 | 2 | 0.04 | 3 |
| 2.25 | 2 | 0.33 | -2 |
| **Independence/Additive** |  |  |  |
| 1425.3 | 2 | < 2.2e-16 | 1421 |
| 11335 | 2 | < 2.2e-16 | 11331 |
| 5566.5 | 2 | < 2.2e-16 | 5563 |
| **Additive/Gender interaction** |  |  |  |
| 10.98 | 1 | 1e-4 | 9 |
| 1.20 | 1 | 0.27 | -1 |
| 6.42 | 1 | 0.01 | 4 |
| **Additive/Coherence interaction** |  |  |  |
| 71.76 | 1 | < 2.2e-16 | 70 |
| 21.73 | 1 | 3 e-6 | 20 |
| 3.55 | 1 | 0.06 | 1 |

Similarly, the independence, additive and the two interaction models were fit using the parametric curves. The independence model was rejected in favor of the additive model for all three tasks (face: *χ*^2^(2) = 1425, p << 0.001; voice: *χ*^2^(2) = 11335, p << 0.001; stimulus: *χ*^2^(2) = 5566, p << 0.001). The gender magnitude interaction described in equation (7) was tested against the additive model and found to be significant for the face and stimulus tasks (face: *χ*^2^(1) = 10.98, p < 0.001; voice: *χ*^2^(1) = 1.2, p = 0.27; stimulus: *χ*^2^(1) = 6.42, p = 0.01). In contrast, an interaction due to the coherence of the stimuli (equation 6) was only significant for face and voice tasks (face: *χ*^2^(1) = 71.8, p << 0.001; voice: *χ*^2^(1) = 21.7, p << 0.001; stimulus: *χ*^2^(1) = 3.55, p = 0.06).

To summarize, there is no evidence that judging “most masculine” vs “most feminine” influences the manner in which observers integrate face and voice gender information whether they are instructed to judge the face, the voice or the stimulus under the additive model. In addition, male and female observers perform similarly, although there is a significant but small trend indicating a sex difference in the contribution of the face component in the voice task. After combining these conditions, the independence model can be rejected for all three tasks, indicating that observers cannot completely suppress the modality that is not attended to. These results suggest that unimodal sensory signals are combined prior to the decision.

Both of the interactions that we tested, an effect of gender magnitude and an effect of intra-stimuli gender congruence, fit some conditions better than a simple additive model. It is somewhat surprising that coherence interactions were not found in the stimulus task suggesting an exclusive congruence between face and voice gender in this task. Interestingly, gender and coherence effects seem to be independent from each other, with the gender effect being non-significant for the voice task and the congruence effect for the stimulus task. This could indicate distinct neural bases, a possibility that is explored further in the imaging experiments.

We examined the accuracy with which each model predicts the observers’ responses on a trial-by-trial basis. For the face task, the accuracy of the additive model increases from 78.96% to 79.22% by including both interactions (+0.26%). For the voice task, the accuracy increases from 83.26% to 83.44% by including both interactions (+0.18%). For the stimulus task, the accuracy increases from 80.49% to 80.58% by adding both interactions (+0.09%). In other words the initial accuracies with the additive models are already high and the improvements in fit, while significant under the likelihood ratio tests and as indicated by the differences in AIC are modest. The improvements, however, mirror the statistical results in that the accuracy is least improved for the stimulus task.

Figure 6A shows schematically an example of how the response predictions for an additive and interaction model differ. For the additive model (dashed lines in the left figure), each line shows the variation of response along the stimulus dimension indicated on the abscissa with the response values on the other dimension added to it. As the models are additive, the displacement of each function is parallel. Here, we have made each of the functions linear for illustration purposes. The interaction model is required when this parallelism is not respected, in the particular example shown here indicated by the varying slopes of the solid lines. However, any other deviation from parallelism is possible. The right graph of Figure 6A shows the difference of the lines from the interaction model and represents the format employed in subsequent analyses of the interaction effects.

**Figure 6:**
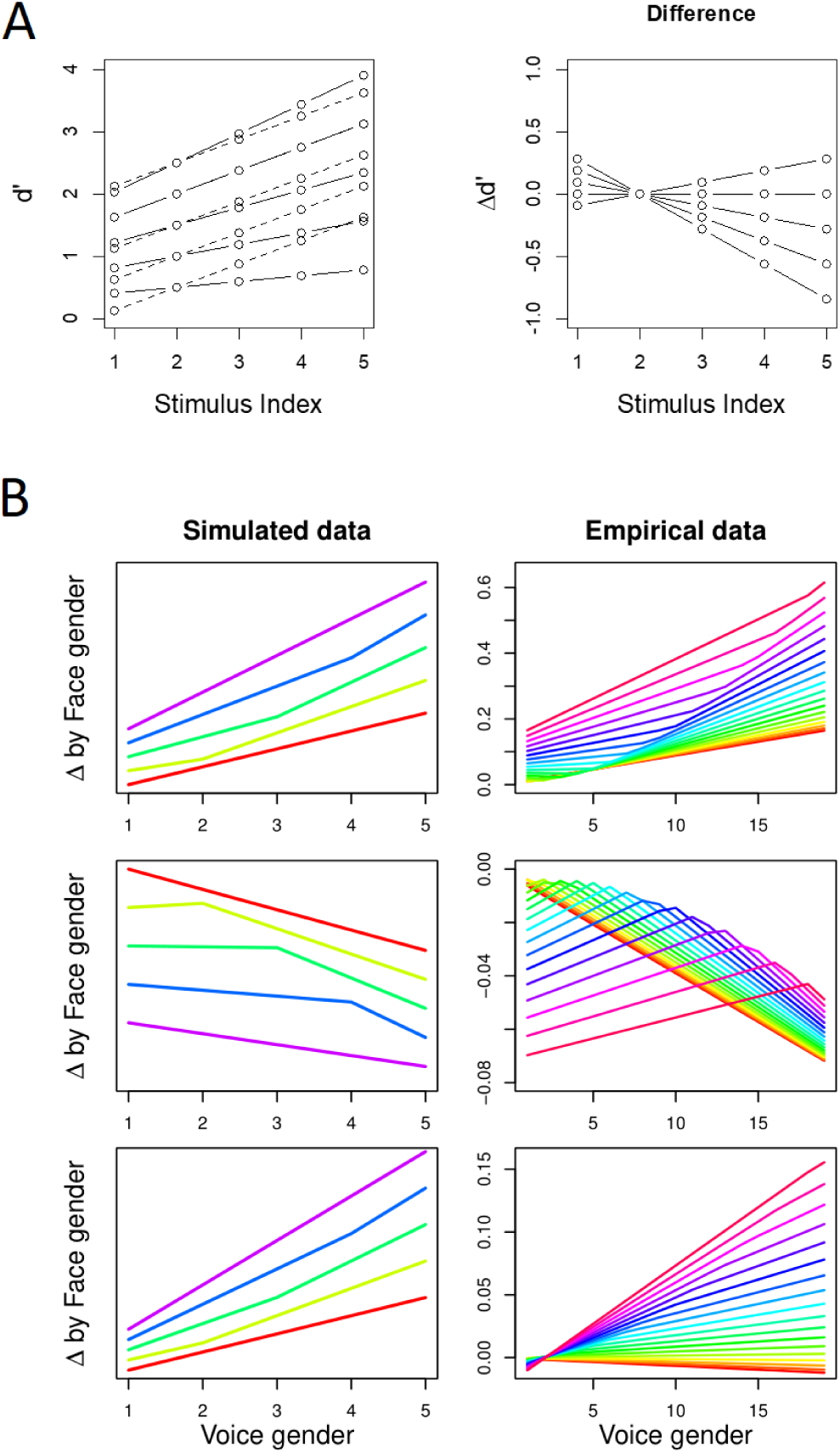
Behavioral interaction effects for gender magnitude and coherence. A. Schematic illustration of the difference in response predictions between additive and interaction MLCM models with arbitrary linear scales. On the left plot, the two models are shown individually for each combination of stimulus index (dashed lines for the additive model, full lines for the interaction model). On the right plot are shown all of the differences between these curves. B. Comparison between interaction and additive models for simulated and empirical data. All plots follow the same format as the right plot from Figure 6A by contrasting interactions models including both a gender coherence and gender magnitude effects with additive modes. On the left side are simulations (with 5 levels for each modality for clarity) in which interaction effects were introduced either by decreasing decision noise in proportion to the distance of each unimodal component from gender-neutral (third row), by increasing decision noise in proportion to the gender distance between each unimodal component (second row), or both (first row). On the right side are the difference between interaction and additive models for our behavioral data (third row: stimulus task, second row: voice task, first row: face task).

In Figure 6B, we compare the interaction models that best fit the three different conditions (right column) with the predicted effects from simulated data including the significant interaction terms. Starting with the bottom set of graphs, the figure on the right shows the difference of interaction and additive fits for the stimulus condition, in which the gender interaction effect was significant. In this case, the greater the gender difference along each axis, the greater its contribution to the interaction. The difference between the additive and interaction fits results in functions that fan out and whose slopes change slightly as the voice gender increases. The lower right graph shows a selection of curves for data simulated to include this type of interaction that display qualitatively similar changes.

The second row of graphs illustrates the coherence interaction, which was significant in the voice condition. In this case, the contribution to the interaction depends on the magnitudes of the difference in gender response between the two modalities for each stimulus. The greater the difference between the face and voice genders, the larger the contribution to the interaction term. The graph on the right shows the difference of additive and interaction fits for the voice condition. The curves rise sharply with voice gender but then break sharply with a negative slope. The breakpoint shifts systematically across face gender values. The graph on the left shows the results from data simulated to include exactly this type of interaction and show qualitatively the same effects of slope change and shifts in the breakpoints.

For the face task, both interactions were found to make a significant contribution to the decision rule. The differences in the additive and interaction models (right panel Figure 6B) show changes in slope with the pattern of change varying across the level of face gender. The simulations of data containing contributions from both such interaction terms display similar qualitative changes in the left panel of figure 6B.

As a whole, Figure 6 illustrates that while the difference between additive and interaction models is much smaller than the main effects (compare ordinate scales between Figures 5 and 6B), the shape of each interaction displays a different signature. The interaction is mostly driven by the gender magnitude effect in the stimulus task (hence the fan-like shape), by the coherence effect in the voice task (hence the inverted U shape) and by a mixture of the two in the face task. Note also that, in the face and the stimulus task, interaction effects increase the contributions compared to the additive model, whereas they have a decreasing effect in the voice task.

In summary, the psychophysical results show that the contributions of each modality vary according to the task by increasing the relevant and attenuating the irrelevant modality. The rejection of the independent model for all tasks means that both face and voice make significant contributions to gender evaluation in all 3 tasks, i.e. there are irrepressible cross-contributions of the voice during face gender evaluation and of the face during voice gender evaluation. Interestingly, the first two graphs of Figure 5 show an asymmetry; the voice contribution is higher in the voice task than the face contribution in the face task and the face contribution is lower in the voice task than the voice contribution in the face task. Finally, we find two independent interaction effects; an effect of gender coherence significant in the face and the voice tasks, and a multiplicative effect of gender magnitude significant in the face and the stimulus tasks, which can be qualitatively compared to simulated results derived by extending the optimal cue combination model under the principles of Signal Detection theory as specified in equations 6 and 7 (see also Supplementary Section 4).

### fMRI results

#### Regions of interest

We selected two areas, the FFA and TVA, activated significantly by our face and voice functional localizers, based on a GLM analysis described in the Material and methods section. Figure 7 illustrates the localization of these areas in one participant; Figures S9 and S10 show the localizations for each individual participant. We computed functional signal-to-noise ratio across these ROIs for responses to the face-voice stimuli used in the main protocol. We found that the mean activity in the TVA was higher than baseline in every condition while the activity in the FFA was higher than baseline during the face and stimulus tasks but not during the voice task (Figure S11). We attribute the large error bars to the small sample size used. We then explored the effects of attentional modulation of response in terms of effective connectivity between FFA and TVA using a DCM analysis.

**Figure 7:**
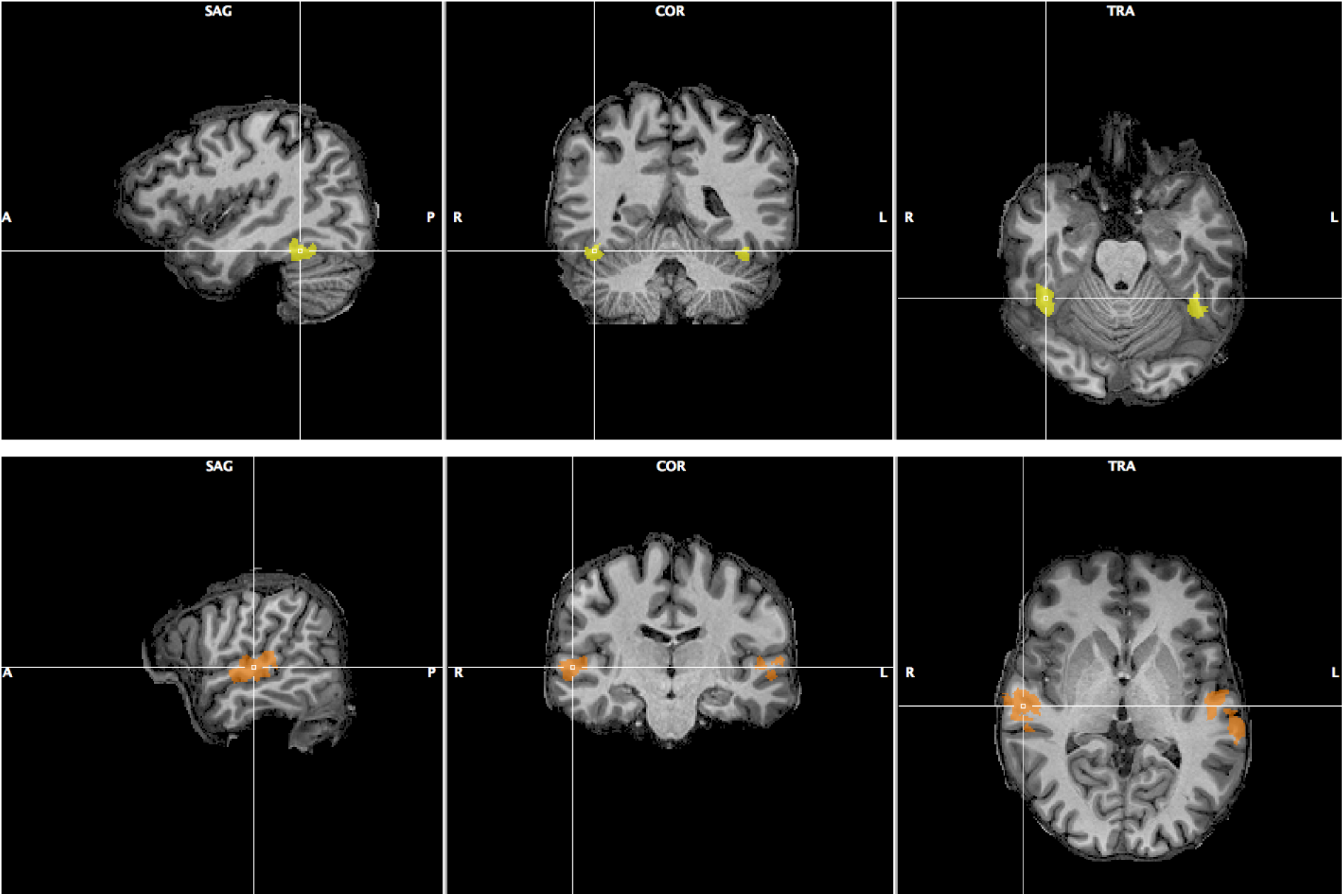
Fusiform Face Area (top row, yellow) and Temporal Voice Area (bottom row, orange) as localized in one of the 12 participants of Experiment 2. Sagittal, coronal and transverse view are shown centered around the right FFA/TVA.

#### Face-voice gender task

So as to engage the attention of a specific modality during imaging sessions in a manner similar to that imposed by the response instructions during the psychophysical tasks, subjects performed a pre-specified gender identification task with respect to either the face, voice or stimulus on randomly signaled trials that were excluded from the subsequent imaging analyses. Prior to the fMRI data analyses, we checked the results of these response trials to evaluate if the participants had correctly performed the tasks. Table 2 shows, for each task performed during the imaging experiments, the mean and standard deviation over 8 subjects of the performance in stimulus identification. There were 40 response trials in total but only 30 were analyzed for each task. In particular, the following trials were excluded from analysis:

- for the face task the 10 trials with a gender neutral face
- for the voice task the 10 trials with a gender neutral voice
- for the stimulus task the 10 trials with incongruent face and voice

The column labeled “Correct” indicates the number of responses that were congruent with the gender of the attended modality (for example in the face task if the masculine sign was on the left, the face was masculine and the observer pressed the left button). “Incorrect” indicates the number of responses that were incongruent with the gender of the attended modality (for example in the previous case if the observer pressed the right button). “Miss” indicates the number of times that the observer did not respond within the 2 second limit. Participants responded in accordance with the attended modality (95%) with less than one incorrect or missed trial per acquisition. Standard deviations are low, indicating little variation between subjects. In summary, the evidence supports that subjects performed the task correctly, which we considered as a validation for the subsequent fMRI data analyses.

**Table 2:** Results of the behavioral task during fMRI recording.

|  | **Correct** | **Incorrect** | **Miss** |
| --- | --- | --- | --- |
| **Face task** |  |  |  |
| mean | 28.50<br>(95%) | 0.75<br>(2.5%) | 0.75<br>(2.5%) |
| sd | 0.93<br>(3.1%) | 0.89<br>(3%) | 0.89<br>(3%) |
| **Voice task** |  |  |  |
| mean | 28.50<br>(95%) | 0.50<br>(1.7%) | 1.00<br>(3.3%) |
| sd | 1.60<br>(5.3%) | 0.53<br>(1.8%) | 1.77<br>(5.9%) |
| **Stimulus task** |  |  |  |
| mean | 26.88<br>(89.6%) | 2.5<br>(8.3%) | 0.62<br>(2.1%) |
| sd | 1.25<br>(4.2%) | 1.20<br>(4%) | 0.74<br>(2.5%) |

#### DCM analysis

Family comparison results are shown in Figure 8, where black and colored bars indicate the exceedance probability of model families, respectively, without and with the arrow corresponding to the model spaces of Figure 4.

**Figure 8:**
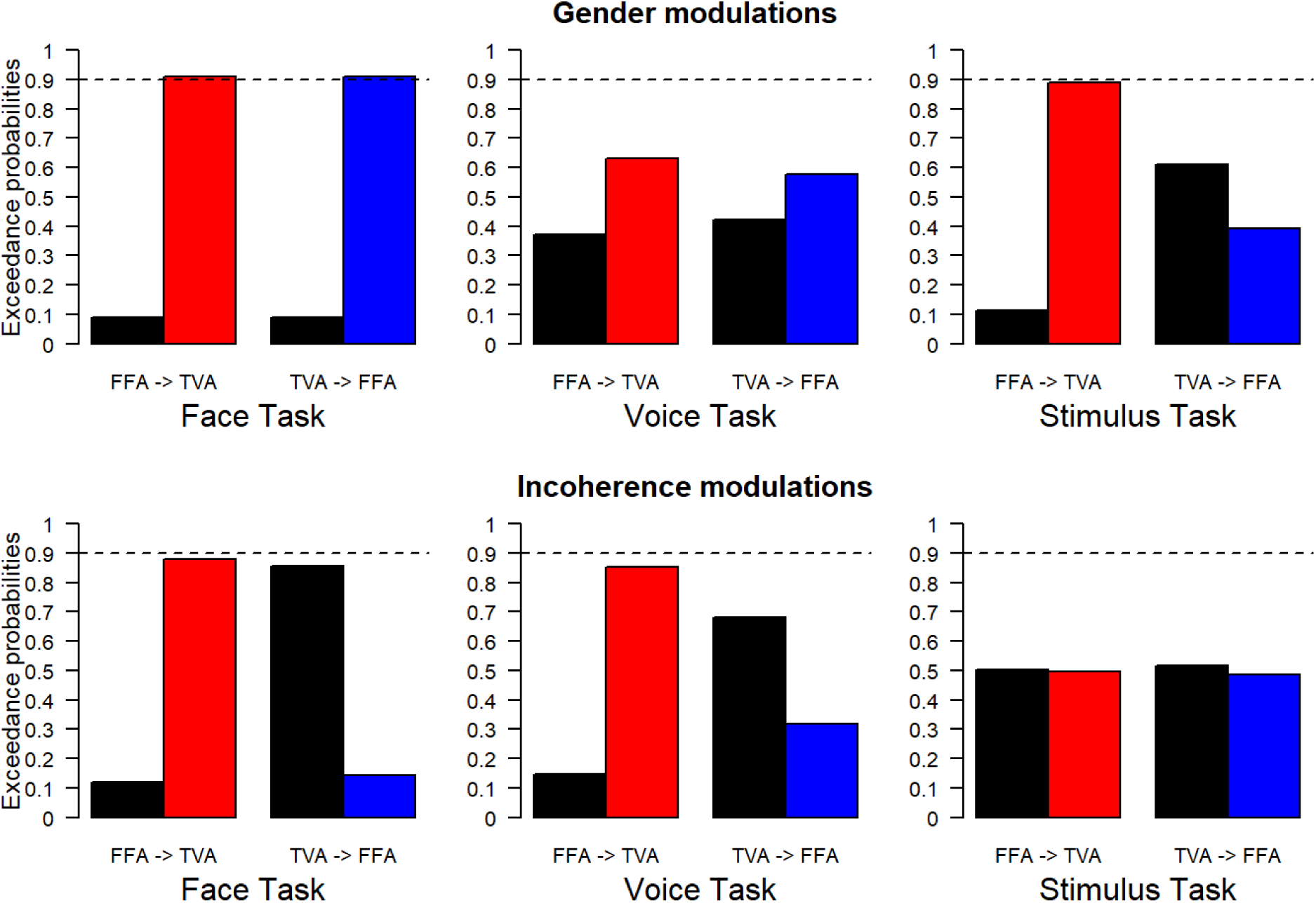
Results of family BMS for all conditions, model spaces and partitions. Top row: modulations of effective connectivity by face/voice gender information. Bottom row: modulations of effective connectivity by face/voice gender incoherence. Within each column, on the left are modulations when subjects are paying attention to face gender, on the middle when they are paying attention to voice gender and on the right when they are paying attention to stimulus gender. Within each graph, on the left are the respective exceedance probabilities for the family of models without modulation from FFA to TVA (in black) versus the family of models with this modulation (in red). On the right are the respective exceedance probabilities for the family of models without modulation from TVA to FFA (in black) versus the family of models with this modulation (in blue). Probability of 0.9 is indicated as a reference for what should be considered strong evidence in favor of a family.

The pattern of results is similar to those obtained from the analyses of the psychophysical data. When subjects’ task was to focus on the gender of the face, there was an interaction effect of gender magnitude in the psychophysics and strong evidence for modulation of effective connectivity from FFA to TVA in response to face gender information (as opposed to gender-neutral faces). A similar pattern was shown in the modulation of effective connectivity from TVA to FFA in response to voice gender information (as opposed to gender-neutral voices). Similarly, there was an interaction of gender coherence in the psychophysical results and also evidence for effective connectivity modulation from FFA to TVA in response to incoherence of gender between the face and the voice. Interestingly, there was also evidence in favor of an absence of modulation (as opposed to simply no evidence in favor) from TVA to FFA in response to gender incoherence.

When the subjects’ task was to attend to the gender of the voice, there was no gender magnitude interaction in the psychophysical analyses and similarly no evidence for effective connectivity modulation in response to gender. There was an interaction of gender coherence in the psychophysical analysis during the face task, the evidence for which was less strong based on p-values and similarly evidence for effective connectivity modulation from FFA to TVA in response to gender incoherence.

When subjects were focusing on the gender of the stimulus, there was evidence of the gender magnitude interaction observed in the psychophysical analyses, and similarly evidence for modulation of the effective connectivity from FFA to TVA in response to face gender information. There was no interaction of gender coherence observed in the psychophysical experiments and similarly no evidence of modulation of effective connectivity in response to gender incoherence

In summary, the results show that changes in effective connectivity parallel the MLCM interaction effects. In addition, in the case of face-voice integration, both the gender magnitude and gender coherence interactions should be considered as early interactions as described in Figure 1 because they correspond to interactions between uni-modal areas. This result was not unexpected given that multimodal effects have been reported as early as primary visual cortex (Petro et al., 2017).

## Discussion

We find that the integration of face and voice information in gender perception can be described by a weighted combination of contributions from each modality and that these weights can be modified by instructing observers to base their judgments on a specific modality or on both. For example, instructing subjects to judge gender on the basis of the face led to an average 5-fold ratio of the face to the voice contributions while instructing subjects to judge gender on the basis of the voice yielded a nearly 8-fold ratio of the voice to face contributions; thus, there is a 40-fold variation in weighting due to the difference in instructions, suggesting that the weights are influenced by top-down processes. Our results confirm those of Watson et al. (2013) by showing that the dominance of a modality is task dependent. The MLCM technique, in addition, enabled us to quantify the specific contributions of each modality. This quantification revealed that the functional dependence on the relative gender of the stimulus of each modality’s response was invariant. In other words, the shape of the curve describing each modality’s contribution did not change with the top-down instruction, suggesting that the contribution of each modality depended on early uni-modal pathways, and the top-down effects could be described as a simple re-weighting of each modality invariant function. Nevertheless, gender decisions could be attributed to a multi-modal integration site, because both modalities contributed significantly to the decisions independently of the task. Such a weighting effect (as opposed to simply increasing the contribution of the relevant modality without affecting the other) supports an optimal use of limited resources.

Comparing the top-down influences of task, there appears to be an auditory dominance. The auditory contribution increases more in the voice task than the visual contribution in the face task, and the visual contribution decreases more in the voice task than the auditory contribution in the face task. Watson et al. (2013) observed the same phenomenon in a gender identification task using nearly an identical stimulus set in which they analyzed the probability of choosing one gender (but see Latinus et al. (2010)). One explanation for the auditory dominance is that there is a greater sexual dimorphism in the auditory than the visual domain for faces; female and male voices differ significantly in fundamental frequency (Puts et al., 2011), and in the stimulus set that we used the voices obviously change in pitch when varying from female to male. Visual changes with gender tend to be distributed across the face and are more difficult to describe (Macke & Wichmann, 2010). In the current stimulus set, our subjective impression was that the differences appear to be related to the sharpness of contours, which is in accordance with previous results linking face gender perception and contrast (Russell, 2009). If the differences in the sensory ranges between modalities were so large, however, we might expect the auditory signals to dominate in the stimulus task and, in fact, they do not; instead both dimensions contribute about equally. Additionally, the response range along the *d’* scale of the visual component in the face task is only about one-third smaller than that for the auditory component in the voice task, suggesting that integration mechanisms can quite easily adjust modality specific weights to compensate for differences in the range of input signals.

Beyond the additive contributions from each modality to gender judgments, we also detected specific and significant interaction effects related to the coherence of gender between modalities and to the magnitude of the gender signal (i.e., differences from neutral gender) within each modality. We simulated these interaction effects in terms of contributions to decision noise based on an optimal cue combination decision rule within a signal detection model and found the simulated results to be in qualitative agreement with the estimates from the data. According to this simulation, when the gender information from one modality is inconsistent with that from the other, the integration process has an increased variance, which leads to a decision bias. For the gender magnitude interaction, the closer the gender within a given modality is to neutral, the larger the variance assigned to it, which also results in decisional biases

The two interaction effects do not occur under the same conditions. Significance of the coherence interaction occurred during the face and the voice tasks; significance of the gender magnitude interaction occurred during the face and stimulus tasks. This led us to explore the interactions between cortical areas implicated in face and voice processing for correlates of the interactions. Using an equivalent face-voice gender categorization task with DCM, we found that modulations of connectivity between the FFA and the TVA mirrored the behavioral interaction effects, consistent with the hypothesis that these effects do not exclusively depend on higher-order multi-modal integration sites.

Mirroring the conditions in which the coherence interaction occurs, the effective connectivity from FFA to TVA was modulated for the face and voice tasks. Such an effect is not observed during the stimulus task, however, perhaps reflecting the reduced weight assigned to the auditory component observed in the psychophysics thereby minimizing its role in integration faced with incoherence of gender across modalities.

Magnitude of gender difference from neutral induced no change in effective connectivity during the voice task but a bilateral modulation during the face task and FFA to TVA modulation during the stimulus task. This parallels the psychophysical results for the gender magnitude interaction that was significant for the face and stimulus tasks but not for the voice task. In the case of the stimulus task, this might represent a mechanism generating a change in weight in both modalities to equalize their contributions to the judgments. Given the default auditory dominance described above in the psychophysics, a similar explanation might apply in the face task. We hypothesize that task-dependent reweighting of the contributions of the modalities might constitute a behavioral correlate of the task-dependent dynamic reorganization of inter-areal hierarchical relations observed in the frequency-specific synchronization of local field potentials in macaque (Bastos et al., 2015).

The two interactions modeled in the psychophysical analyses correspond to the integration of information discrepancies that must be integrated in order to make a judgment in the specific task. In the case of the coherence interaction, the voice and face gender information is in conflict with respect to the gender identity of the stimulus. On the other hand, the gender magnitude interaction depends on the strength of the gender signal and, thus, could be related to the precision of its encoding. These differences could be related to the multiplicity of feedback pathways (Markov et al., 2014) having distinct functional roles (Bergmann et al., 2019; Markov & Kennedy, 2013; Shipp, 2016).

One computational theory about the role feedback signals is that they contribute to the construction of Generative models of the outside world (Friston, 2002, 2005; Mumford, 1992; Rao & Ballard, 1999). In this framework each processing step should be conceived as trying to predict its feedforward inputs using the feedback signals it receives. Predictions (which can be interpreted as some internal representation, such as a template) are transmitted to lower levels via feedback signals, and prediction errors (which can be interpreted as some function of model residuals) are transmitted to upper levels via feedforward signals. In this context a conflict in the integration of two signals would lead to an error signal propagated from lower to higher areas. If this is what the gender incoherence modulation reflects, then it places the FFA at a lower hierarchical level than the TVA.

Anatomically, the FFA and TVA belong to streams of two different modalities, making their hierarchical relations indirect. Support for the hypothesis that the TVA is hierarchically higher than the FFA can be found in the macaque where a face responsive patch comparable to the FFA is situated in the TEpd area (Lafer-Sousa and Conway, 2013) and voice responsive neurons that have been argued to form a TVA-like patch (Perrodin et al., 2011; Belin et al., 2018) are situated in the parabelt area. Analysis of anatomically derived measures of hierarchy that are based on laminar connectivity patterns (Markov et al., 2014) indicate that the TEpd is indeed hierarchically lower than the parabelt area PBr. Note, however, that inter-species homologies must be applied with caution, and effective connectivity does not necessarily imply monosynaptic connectivity.

We have demonstrated that top-down influences can dramatically change the weight of visual and auditory contributions in a simple gender perception task. In addition, we found evidence for specific interactions in the decision processes that reflect both the coherence of the gender information across modalities and the magnitude of the gender information. The conditions that generated these interactions were accompanied by specific changes in effective connectivity between cortical areas implicated in face and voice processing. The results support a role for multiple feedback processes in multi-modal integration.

## Supporting information

Supplementary sections

## Acknowledgment

We thank Michel Dojat for critical comments and discussion of DCM, Franck Lamberton and Danielle Ibarrola for technical assistance in data acquisition, Dr Alain Nicolas for medical monitoring of the subjects, Pascal Belin and Rebecca Watson for the face-voice stimulus set.

## Author Contribution

Project conception HK KK; Procured funding HK; Experimental design CA, PG, KK; Data collection: CA, PG; Data analysis CA, PG, KB, KK; Wrote the paper CA, PG, HK, KK; All authors revised successive versions of the paper.

