## Supplementary sections for "Voice and Face Gender Perception engages multimodal integration via multiple feedback pathways"

**Supplementary section 1**

Previous MLCM experiments used a stimulus set with a small number of levels for each dimension and tested all combinations of these levels for a given number of repetitions. For example Ho, Landy, and Maloney (2008) used 5 levels along 2 dimensions, yielding 5x5 = 25 combinations and 25x24/2 = 300 unique, unordered pairs that they repeated three times, giving 900 stimuli per observer. In this experiment we used a stimulus set with 18 levels of morphing for the face and 19 for the voice, yielding 19x18 = 342 combinations and 342x341/2 = 58311 unique, unordered pairs. Considering a trial duration of approximately 2 seconds, it would take over 30 hours for a participant to respond to all pairs once.

Using simulation, Knoblauch and Maloney (2012) proposed that equivalent levels of precision for the estimates could be achieved by either sufficiently sampling randomly from a larger stimulus set or testing all combinations of a smaller stimulus set. In their simulation, they examined the influence of the number of levels and the percentage of sub-sampled pairs on the expected variability of the estimates. Here, we perform similar simulations but examine instead the consistency of the estimates, i.e., how close they are to the ground truth of the underlying response function.

Description of simulation: We used a pair of fixed internal response functions as ground truth to generate all simulated data based on power law fits to a data set of one of the observers from the Ho et al. (2008) study who judged which of a pair of surfaces is bumpier for two surfaces covarying independently in roughness and glossiness in an MLCM experiment. These data are available as the BumpyGlossy data set in the MCLM package (https://CRAN.R-project.org/package=MLCM, Knoblauch and Maloney (2014)). Figure S1 shows the estimated Bumpy (white disks) and Glossy (black disks) additive contributions for each component obtained by maximum likelihood estimation using the function *mlcm* in the MLCM package. The ground truth functions were obtained by fitting an additive MLCM model to these data by maximum likelihood using the formula interface with the additive response constrained to be a weighted sum of power functions, $w_1 B^{e_1} + w_2 G^{e_2}$, where *B* and *G* refer to the bumpy and glossy internal responses, respectively. The best fitting estimates of the parameters were *w* = (1.7, 0.1) and $e$ = (0.8, 1.4) and the fits are shown as solid and dashed curves for Bumpy and Glossy contributions, respectively, in Figure S1. We used these response functions as the true underlying responses and generated stimulus sets with 5, 6, 8, 10 and 20 levels along each dimension, resulting, when both dimensions were paired, in stimulus sets of 25, 36, 64, 100 and 400 stimuli, respectively. The number of pairs for a complete set of trials then was 300, 630, 2016, 4950, 79800, respectively. For each stimulus size, five repeated complete sets of stimulus pairs were generated to simulate an experiment.

For each size of stimulus set, 1000 simulated experiments were run for the complete set repeated 5 times. On each simulated trial, the response difference to the pair of stimuli was computed based on the response functions in Figure S1 to which was added Gaussian noise with mean zero and variance equal to one. If the noise contaminated response was greater than 0, then the first stimulus in the pair was chosen, otherwise the second. The values of the contribution scales for each experiment were then estimated by maximum likelihood as above. The root mean square error was then calculated between the simulated values and the ground truth response functions.

The red symbols in Figure S2 show the results for the complete data sets plotted as a function of the number of stimulus pairs tested on double logarithmic coordinates. The consistency of the estimates improves linearly as a function of the number of samples plotted in this fashion and falls along a line of slope -0.5, indicating an inverse square root law.

We then repeated the simulations but instead of running all pairs, we subsampled 10% and 50% of the complete data sets randomly. The results are shown in white and blue, respectively. All of the points except those for the 5x5 and 6x6 data sets sampled at 10% fall along or close to the inverse square root law. The different sampling rates are represented by color differences while the full stimulus set by different shapes. The results demonstrate a significant range over which stimulus set size and subsampling rate trade-off to produce equivalent levels of consistency in the estimates. Only for small stimulus sets and low sampling rates does the consistency deviate from the inverse square root law. For comparison to our situation, we also simulated 3% subsampling from a 20x20 stimulus set. The results are shown as the orange inverted triangle and generate results similar to 5x5 data set at 50% sampling. Thus, in our case, by subsampling 1500 pairs from an 18 by 19 set, we should expect to obtain a precision comparable to subsampling 150 pairs from a 5 by 5 stimulus set.

To test this equivalence, one subject (author CA, note that this preliminary data was not used for the main experiment) performed 4 conditions of the experiment (judging the most masculine face, the most masculine voice, the most feminine face and the most feminine voice) both with the 18x19 set (subsampling 1500 trials) and with a 5x5 set (exhausting it 5 times for 1500 trials). Figure S3 shows the comparison between the MLCM estimates from the two experiments. The white disks indicate the results of subsampling the larger set and the filled disks (blue for face judgments, red for voice) those for the exhausted smaller set. The judgments of which was more feminine result in functions that were reflected about the abscissa.

The error bars are 95% confidence intervals on the 5 repeated sessions. The curves were obtained by fitting the responses with a Generalized Additive Model (Wood, 2017). The shaded envelopes show 95% confidence intervals about the fitted curves. There were no significant differences in any condition between the results from the two sets, thereby supporting the equivalence of the procedures.

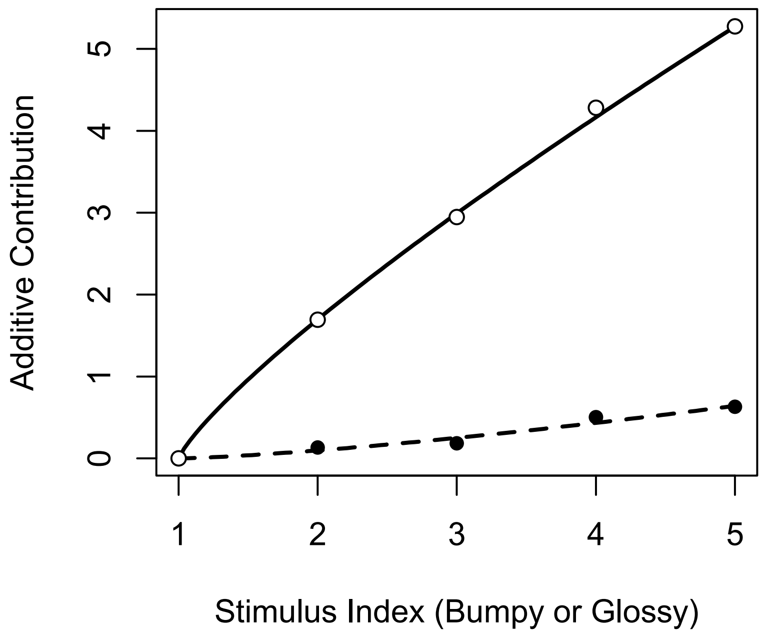


**Figure S1. Estimated Bumpy (white disks) and Glossy (black disks) additive contributions for each component in the BumpyGlossy dataset of the MLCM package obtained by maximum likelihood estimation. Solid and dashed curves are ground truth functions estimated for Bumpy and Glossy contributions, respectively.**

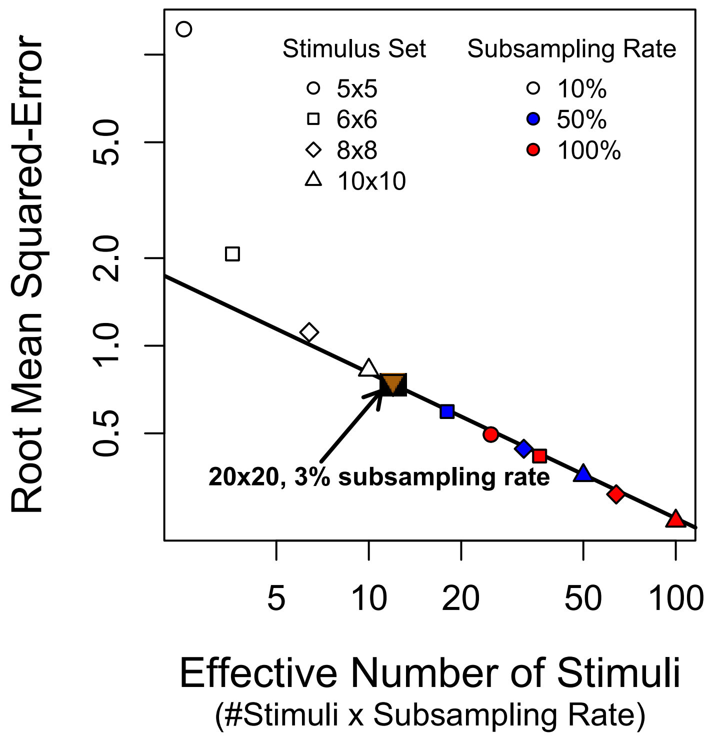


**Figure S2. Accuracy of simulated MLCM results given several parameters. Accuracy is represented in ordinates as the root mean squared-error between 1000 simulated experiments and ground truth, and as a function of the effective number of sampled stimuli in the experiment in abscissa (double logarithmic scales). Symbols represent different sizes of stimulus sets and colors different rates of random subsampling within these sets. The line represents a linear improvement of the estimates as a function of the number of samples, indicating an inverse square root law.**

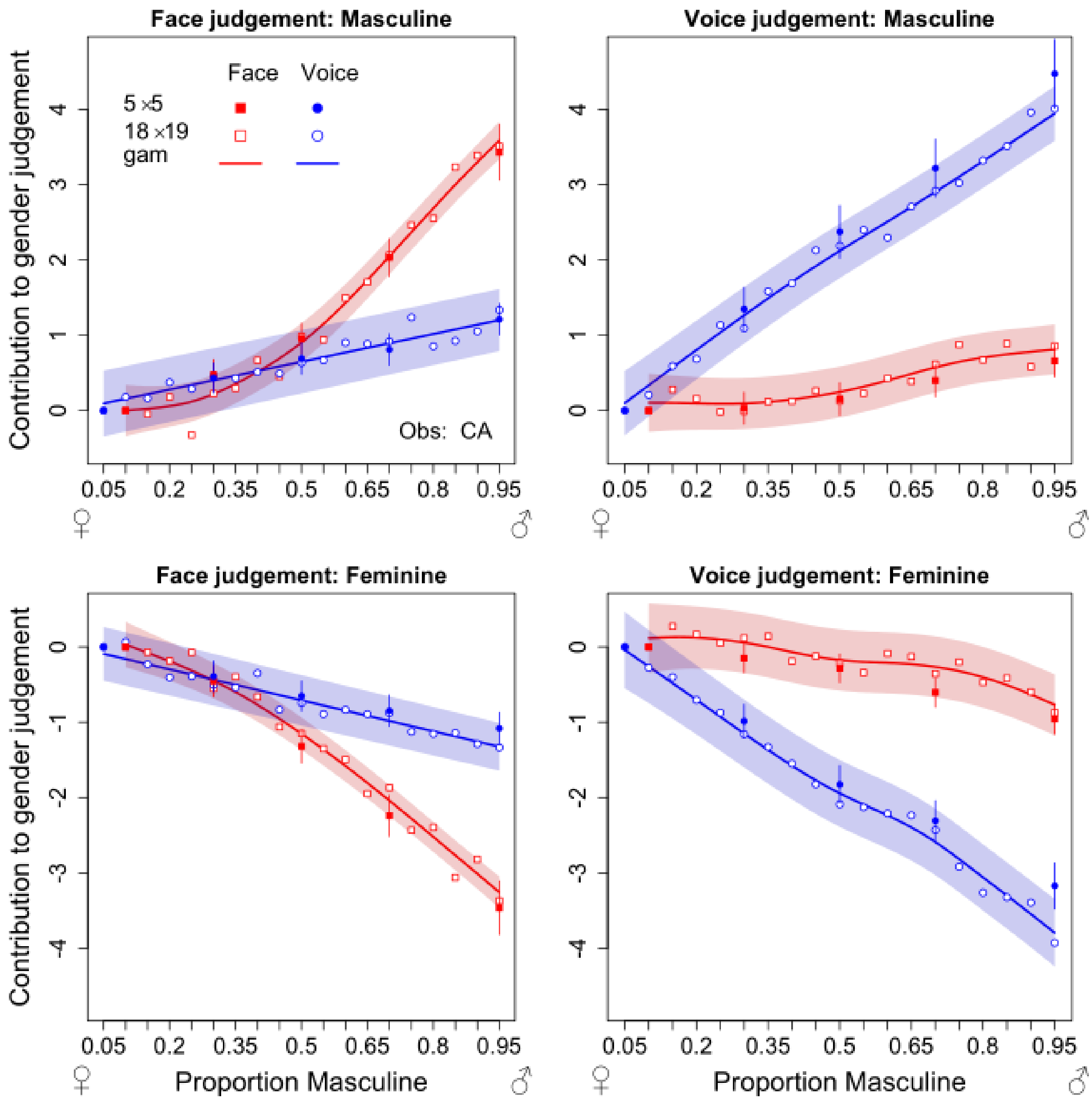


**Figure S3. Contribution of the masculinity of face (in red) and voice (in blue) while an observer judged the most masculine or feminine face or voice in a face-voice MLCM experiment. Filled disks were obtained using a 5x5 stimulus set (exhausting it 5 times for 1500 trials), while white disks were obtained using an 18x19 set (subsampling 1500 trials). The error bars and shaded areas represent 95% confidence intervals and the curves were obtained by fitting the responses of the 18x19 set with a Generalized Additive Model.**

**Supplementary section 2**

Controls for the behavioral experiment. Figures S4 to S6 present individual results of the Generalized Linear Mixed-effects Model (GLMM) for all 36 subjects arranged by condition. Figure S7 contrasts the aggregated auditory and visual contributions of female and male participants in the voice task for the non-parametric GLM model, under which they were found to significantly differ. Figure S8 shows the average values for the visual (left) and auditory (right) components for each of the three tasks, normalized to a common ordinate scale, and normalized shapes approximated with a non-linear least square approach.

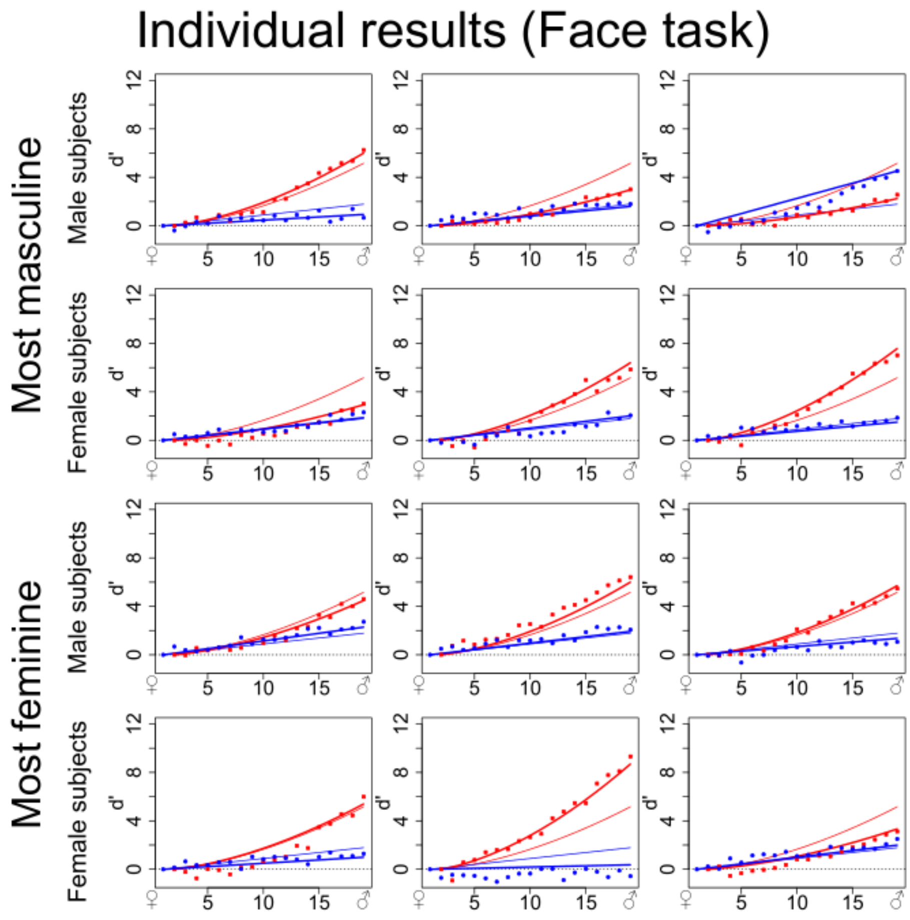


**Figure S4. Individual results from 12 observers doing the Face task in experiment 1. They are regrouped by condition ("most masculine" and "most feminine", in which case contributions are reversed) and by gender. On the**

abscissa are gender morphing levels for both the face and the voice from 1 to 19 (feminine to masculine), in ordinates contributions in d’ units (red: face, blue: voice). Points: non-parametric model. Thin lines: fixed effects of mixed parametric model. Thick lines: individual random effects.

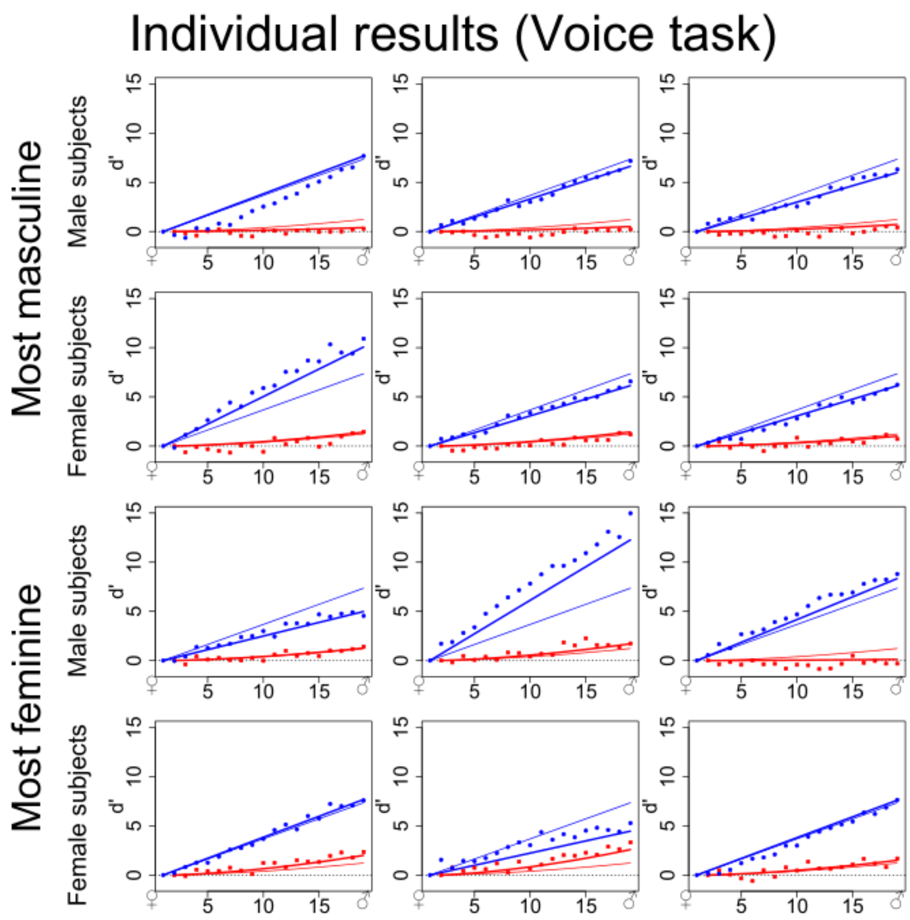


**Figure S5. Individual results from 12 observers doing the Voice task in experiment 1. They are regrouped by condition ("most masculine" and "most feminine", in which case contributions are reversed) and by gender. On the abscissa are gender morphing levels for both the face and the voice from 1 to 19 (feminine to masculine), in ordinates contributions in d’ units (red: face, blue: voice). Points: non-parametric model. Thin lines: fixed effects of mixed parametric model. Thick lines: individual random effects.**

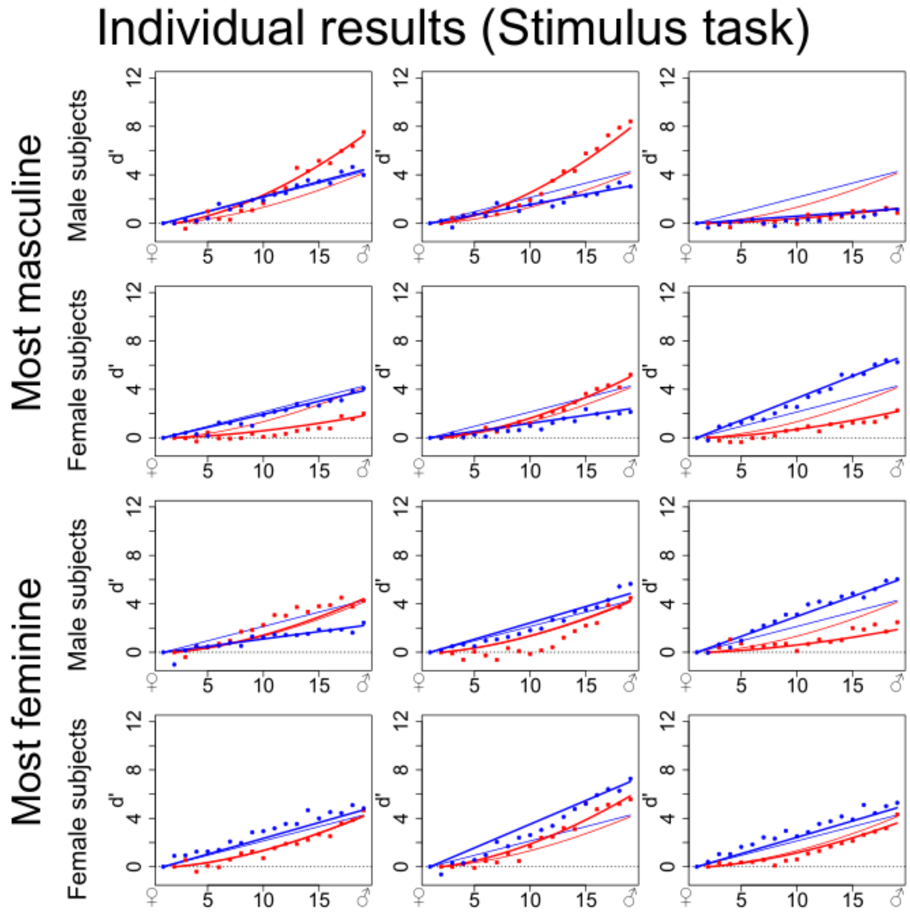


**Figure S6. Individual results from 12 observers doing the Stimulus task in experiment 1. They are regrouped by condition ("most masculine" and "most feminine", in which case contributions are reversed) and by gender. On the abscissa are gender morphing levels for both the face and the voice from 1 to 19 (feminine to masculine), in ordinates contributions in d' units (red: face, blue: voice). Points: non-parametric model. Thin lines: fixed effects of mixed parametric model. Thick lines: individual random effects.**

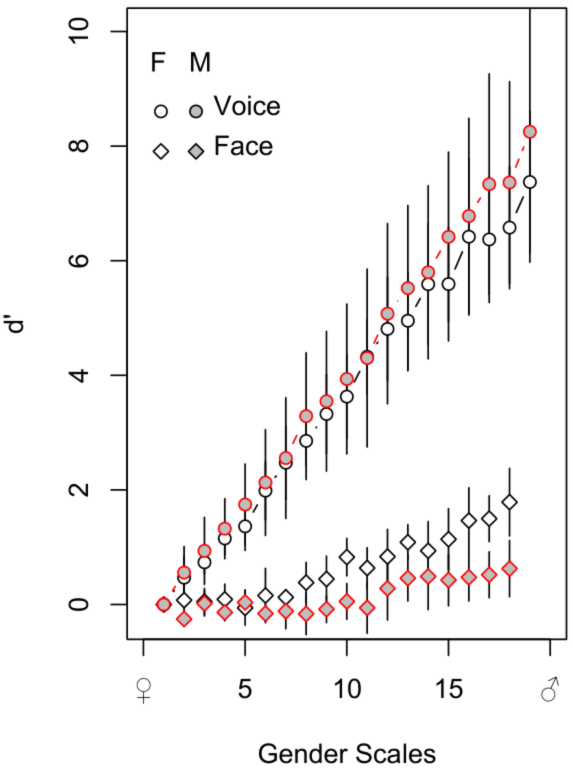


**Figure S7. Contrast of the aggregated auditory and visual contributions (circle and diamond points, respectively) of female and male participants (blank and shaded points, respectively) performing the voice gender comparison task.**

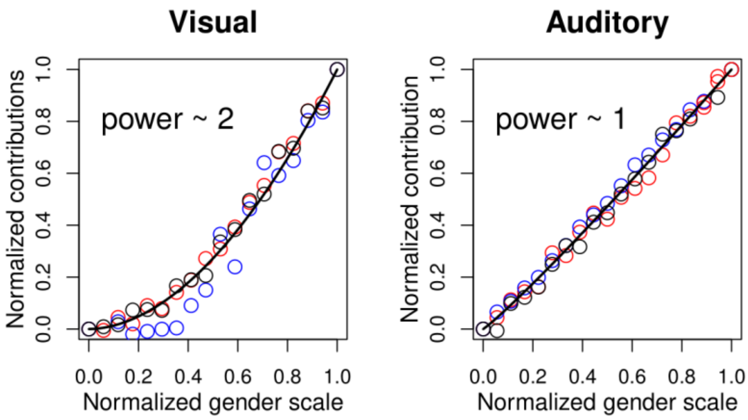


**Figure S8: Non-linear least square approach used to approximate the shape of visual (left) and auditory (right) contributions from mean estimates of the additive non-parametric model applied to all subjects. The points are the mean estimates as a function of normalized morphing level for each task: face (red),**

**voice (blue) and stimulus (black), normalized so that the maximum value is at 1. Data from "most masculine" and "most feminine" conditions are aggregated after inverting responses.**

**Supplementary section 3.**

Controls for the fMRI experiment.

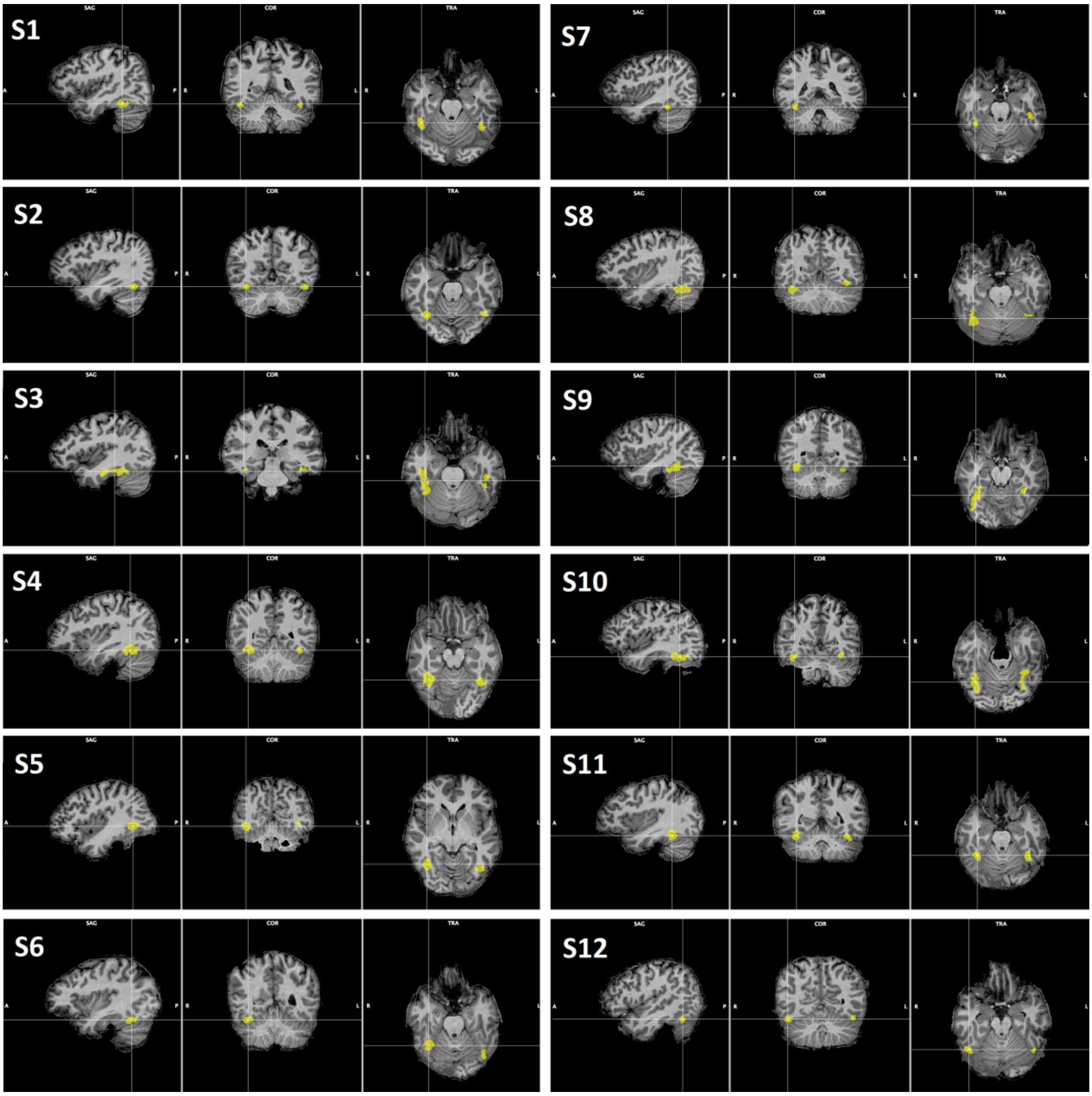


**Figure S9. Fusiform Face Area as localized in each of the 12 participants. Sagittal, coronal and transverse view are shown centered around the right FFA.**

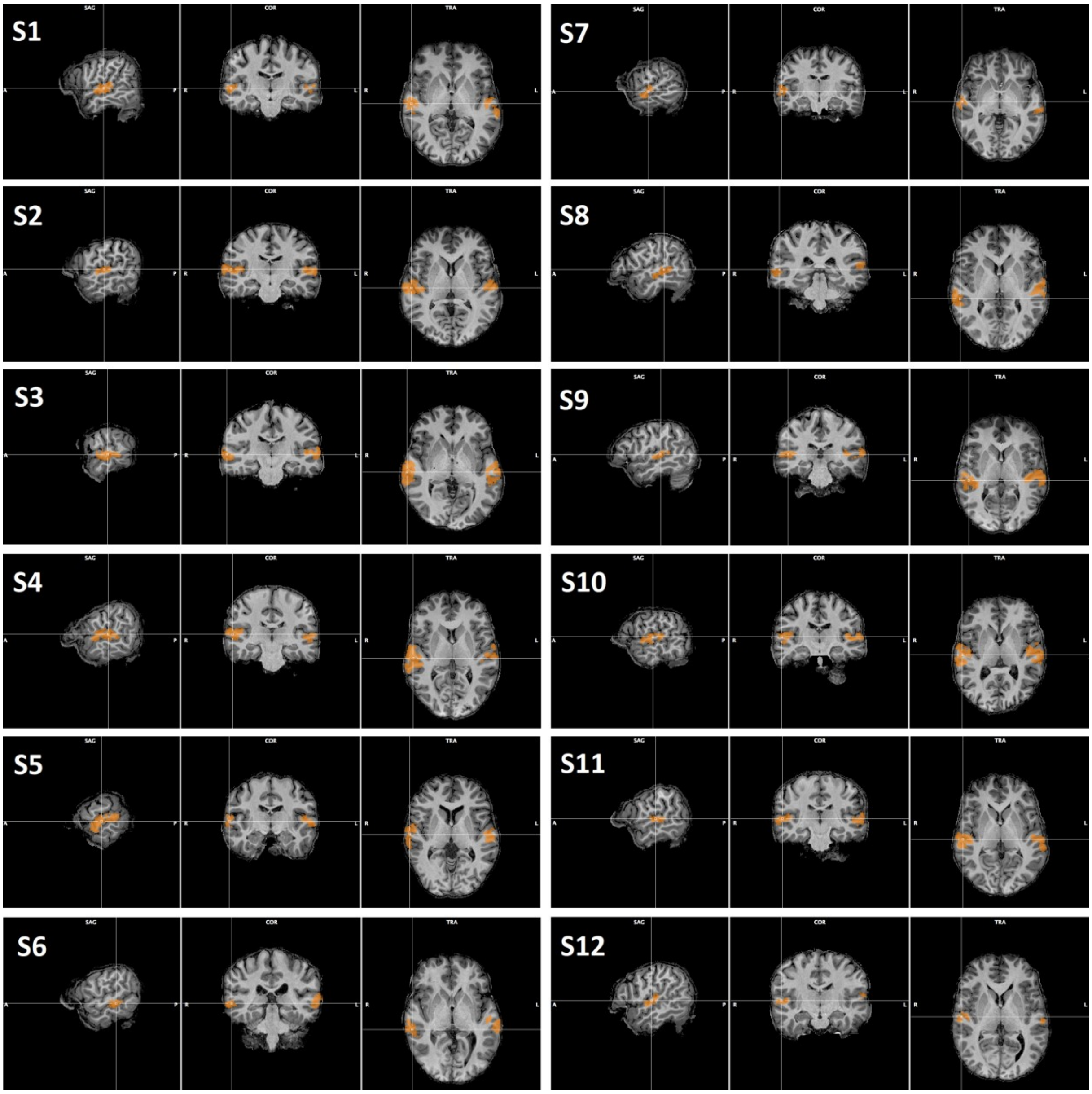


**Figure S10. Temporal Voice Area as localized in each of the 12 participants. Sagittal, coronal and transverse view are shown centered around the right TVA.**

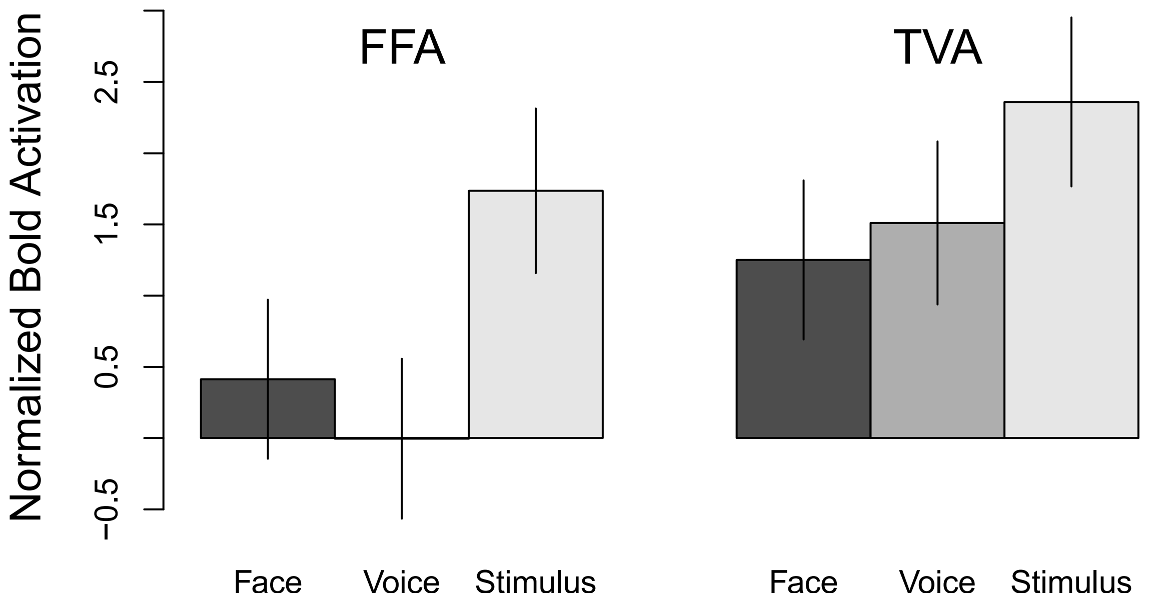


**Figure S11. Initial GLM analysis of the FFA and TVA ROI's. Twelve subjects were exposed to the same sets of multimodal stimuli, faces and voices but participated in one of three conditions at a time on which they were instructed to judge the gender of the face, voice or stimulus on a subset of control trials (indicated on the abscissa). The BOLD signals from these trials were not included in the analyses. Each subject participated in 2 of the 3 conditions on different days; yielding 8 subjects per condition. For each subject, the condition was analyzed with respect to a baseline, defined as the average signal for each voxel over the run. The GLM was run for contrasts in the two ROI's, FFA and TVA. The beta values were adjusted by their standard errors and the data were analyzed with a linear mixed-effects model (Bates, Mächler, Bolker, & Walker, 2015). The error bars are standard errors estimated by a bootstrap method.**

**Supplementary section 4**

When the same stimulus is presented repeatedly to an observer (without any habituation effect), and the observer is asked to evaluate it according to some specific feature, e.g. in the current case, its perceived masculinity, we assume that there is some variability in the observer's responses due to ambiguity in the stimulus and/or stochastic fluctuations in the observer's internal response. Signal Detection Theory (Green & Swets, 1966; Macmillan & Creelman, 2004) hypothesizes that, over many iterations, the decision process can be modeled by a random distribution that we assume, for simplicity, is Gaussian:

$$\psi_{\mathrm{S}}(\phi_{\mathrm{S}}) \sim \mathcal{N}(\mu_{\mathrm{S}}, \sigma^2)$$

where $\phi_S$ is the physical level of the stimulus for the observed feature, $\mu_{\mathrm{S}}$ the mean perceived level (that depends on the morphing level or percent mixing between masculine and feminine reference stimuli) and $\psi_{\mathrm{S}}$ the perceived level of masculinity of the stimulus. See also Figure S12.

Consider a stimulus composed of two independent sources of gender information: visual face, V, and auditory voice, A, components. To evaluate the masculinity of such a stimulus, the observer must combine these two cues in some fashion into a single gender percept.

Clark and Yuille (2013) defined two classes of models to describe how several cues can be fused into a single percept. In weak fusion models each cue is first processed separately in distinct and independent modules, after which only a given rule is used to combine them. By contrast, strong fusion models de-emphasize modularity and allow interaction effects at any processing level. Young, Landy, and Maloney (1993) and (Landy, Maloney, Johnston, & Young, 1995) established the modified weak observer model as a variant of weak fusion that includes early interactions to make cues commensurable (i.e. $\psi_{\mathrm{V}}$ and $\psi_{\mathrm{A}}$ are expressed in the same units on the same perceptual axis and can be directly added), other potential interaction effects when the discrepancy between individual cues is beyond that typically found in natural scenes (i.e., involves choosing $\phi_{\mathrm{V}}$ and $\phi_{\mathrm{A}}$ such that $\psi_{\mathrm{V}} \approx \psi_{\mathrm{A}}$ to prevent such effects) and a weighted average cue combination rule with weights dynamically chosen to minimize variance of the final estimate (which in our case means that $\psi_S = \psi_V \cdot w_V + \psi_A \cdot w_A$ with $w_V + w_A = 1$ with $w_V, w_A$ chosen to minimize $\sigma_S^2$).

The above described model can be formalized as:

$$\psi_S \sim \mathcal{N}(\mu_V, \sigma_V^2) \cdot w_V + \mathcal{N}(\mu_A, \sigma_A^2) \cdot w_A = \mathcal{N}(\mu_V \cdot w_V + \mu_A \cdot w_A, \sigma_V^2 \cdot w_V + \sigma_A^2 \cdot w_A)$$

To minimize the variance of the final estimate, weights are chosen such that $w_V \propto \sigma_V^{-2}$ and $w_A \propto \sigma_A^{-2}$ (Oruç, Maloney, & Landy, 2003):

$$\psi_S \frac{\mathcal{N}(\mu_V, \sigma_V^2)}{\sigma_V^2} + \frac{\mathcal{N}(\mu_A, \sigma_A^2)}{\sigma_A^2} = \mathcal{N}\left(\frac{\mu_V}{\sigma_V^2} + \frac{\mu_A}{\sigma_A^2}, \frac{1}{\sigma_V^2} + \frac{1}{\sigma_A^2}\right)$$

Adding the constraint that $w_V + w_A = 1$, we obtain $w_V = \frac{\sigma_V^{-2}}{\sigma_V^{-2} + \sigma_A^{-2}}$ and $w_A = \frac{\sigma_A^{-2}}{\sigma_V^{-2} + \sigma_A^{-2}}$:

$$\psi_S = \mathcal{N}\left(\frac{\mu_V \cdot \sigma_V^{-2} + \mu_A \cdot \sigma_A^{-2}}{\sigma_V^{-2} + \sigma_A^{-2}}, \frac{\sigma_V^2 \cdot \sigma_A^2}{\sigma_V^2 + \sigma_A^2}\right)$$

The consequences of this model are illustrated in Figure S13: increasing either variance brings the variance and the mean of the combined percept closer to the more reliable estimate. The combined percept is always expected to be more precise than either isolated unimodal estimate. Note that, the observer is predicted to make binary decisions about combined stimuli (i.e. classify them as either feminine or masculine) in proportion to the position of its density distribution with respect to a gender-neutral reference (left or right, respectively).

Observers using this maximization rule with respect to reliability are called statistically optimal and this type of response has been empirically verified in a number of domains for human multimodal integration (Alais & Burr, 2004; Ernst & Banks, 2002; Hartcher-O'Brien, Di Luca, & Ernst, 2014).

As already mentioned, one limitation of this model is its dependency on the condition $\psi_V \approx \psi_A$. In other words complementary interaction effects are expected to come into play in the case of inconsistencies beyond the discrimination threshold between the cues. Figure S14 illustrates that, in the absence of such interaction effects, the resulting percept is predicted to be the same when combining visual and auditory cues as long as the variances and $\frac{\psi_V + \psi_A}{2}$ remain constant.

Figure S15 shows an extension of the model including an interaction effect of coherence. When, for example, one modality is very masculine and the other very feminine, we make the hypothesis that the overall stimulus will be perceived as less reliable than in a more coherent context. Consequently we increase the variance of each unimodal estimate in proportion to its distance before applying the optimal integration scheme. This has the effect of lowering also the variance of the combined estimate, which biases the decisions because a higher proportion of the distribution falls on the left side of the gender neutral reference.

Beyond this effect of coherence, it is also conceivable that there is an effect of the quality of gender information contained in each modality, which is proportional to its magnitude compared to gender-neutral. For example, a face that is very clearly masculine might engender a more precise representation than a more gender-neutral face. Such effects would become more significant with greater incoherence between cues due to an increase in the difference in precision, and once again the resulting change in variance of the final estimate would bias the decisions. An implementation is shown in Figure S16.

In MLCM models, non-additive combination terms are usually introduced as non-specific interaction effects, but we can instead parameterize them according to a priori hypotheses. In our case we define a model introducing an internal coherence parameter and/or an effect of gender magnitude to take into account the quality of gender information.

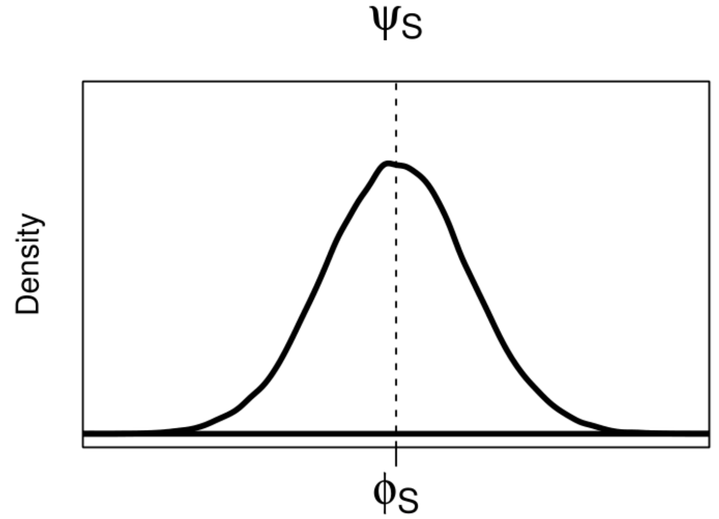

**Figure S12.  Hypothetical distribution of perceived masculinity of stimulus S over many trials with Gaussian distributed error.**

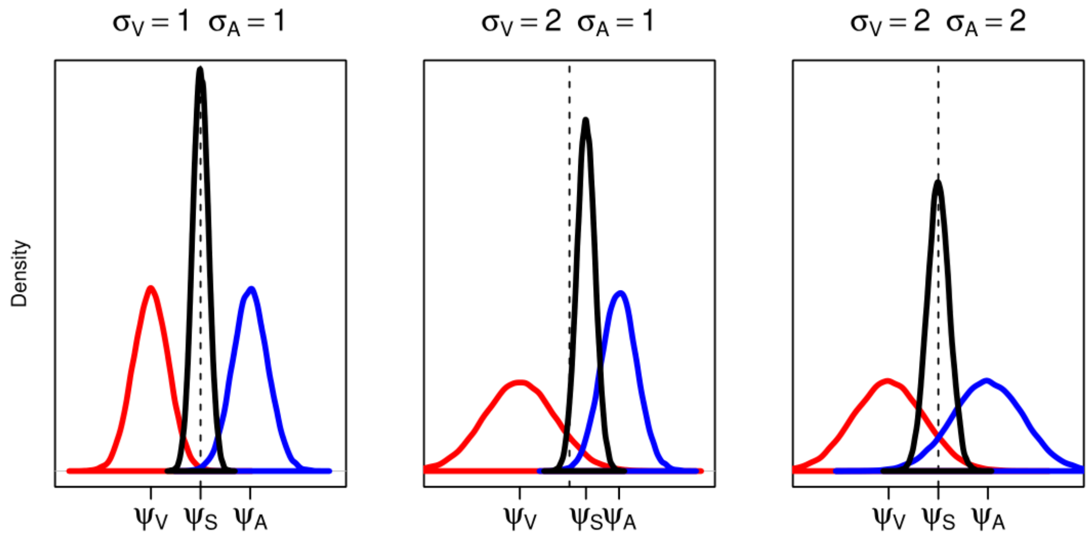


**Figure S13.  Optimal cue combination with varying noise. Visual and auditory unimodal estimates are reported on the same axis (in red and blue respectively), as is the combined percept (in black). The vertical dotted line represents a gender neutral reference, whith the visual estimate being feminine and the auditory estimate masculine. Left: when both variances are equal the combined percept is at the middle with a lower variance than both. Middle: when one variance goes up, the combined percept gets closer to the other modality and its variance also goes up. Right: when both variances go up, the combined percept stays at the middle but with a lower precision.**

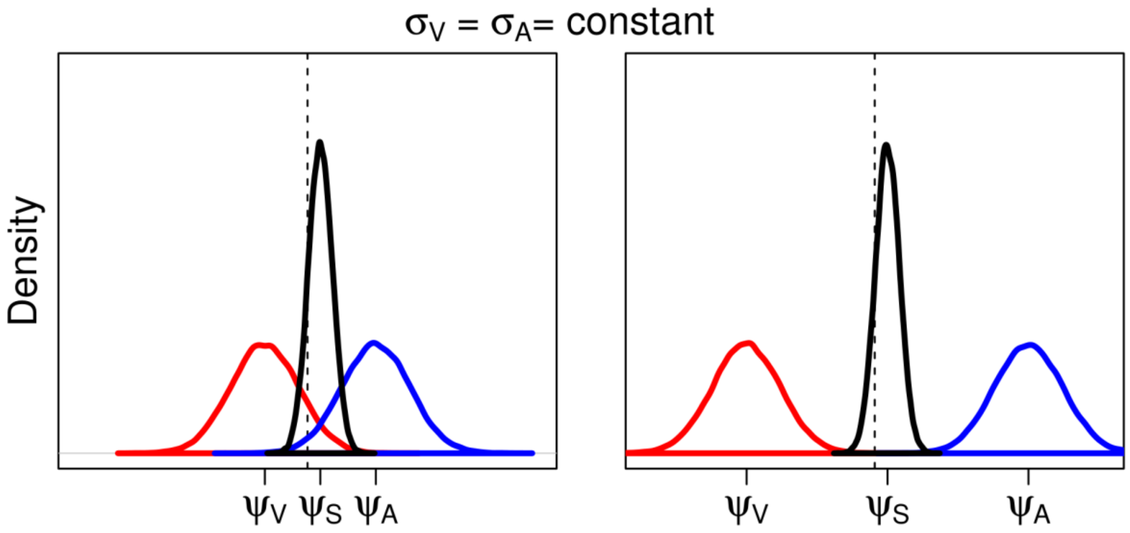


**Figure S14. Optimal cue combination with low (left) and high (right) perceptual difference between modalities in the absence of interaction effects (constant variances). Visual, auditory and combined estimates are reported in red, blue and black respectively. The vertical dotted line represents a gender neutral reference, with the visual estimate being feminine and the auditory estimate masculine. The combined distribution is shifted towards the less gender-neutral auditory signal. It remains the same across plots.**

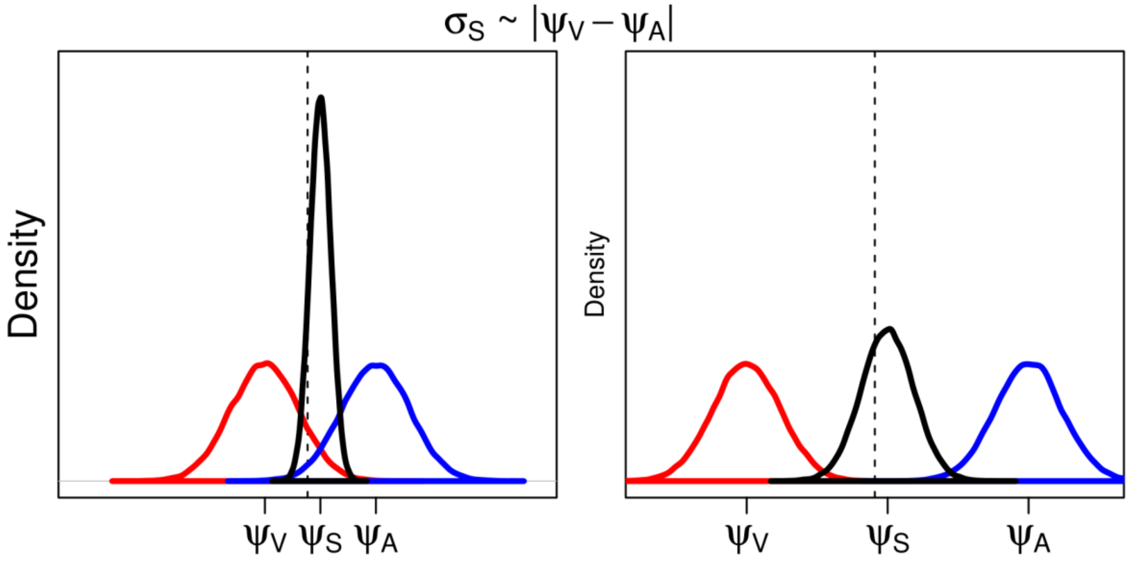


**Figure S15. Optimal cue combination with low (left) and high (right) perceptual difference between modalities and an interaction effect of gender coherence (lower variance for stimuli further apart). Visual, auditory and combined**

estimates are reported in red, blue and black respectively. The vertical dotted line represents a gender neutral reference, whith the visual estimate being feminine and the auditory estimate masculine. The combined distribution is shifted towards the less gender-neutral auditory signal. Its variance is lower on the left side, biasing the decision further.

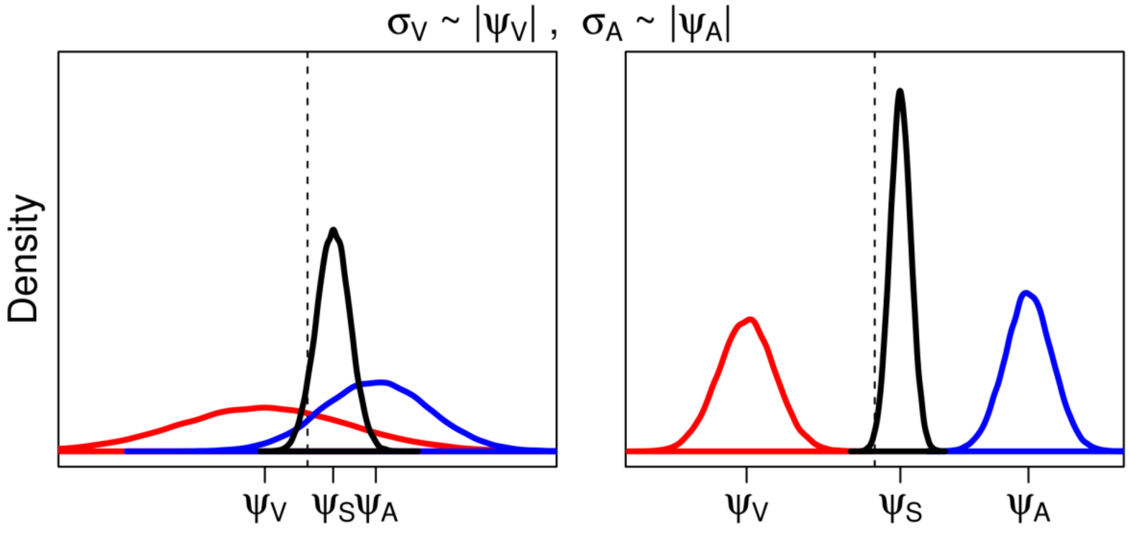


**Figure S16.** Optimal cue combination with low (left) and high (right) perceptual difference between modalities and an interaction effect of gender information (higher variance for stimuli closer to gender-neutral). Visual, auditory and combined estimates are reported in red, blue and black respectively. The vertical dotted line represents a gender neutral reference, whith the visual estimate being feminine and the auditory estimate masculine. The combined distribution is shifted towards the less gender-neutral auditory signal. Its variance is lower on the right side, biasing the decision further.
